# XopM, a FFAT motif containing type-III effector protein from *Xanthomonas*, suppresses PTI responses at the plant plasma membrane

**DOI:** 10.1101/2024.03.06.583702

**Authors:** Charlotte Brinkmann, Jennifer Bortlik, Margot Raffeiner, Suayib Üstün, Frederik Börnke

**Affiliations:** Plant Metabolism Group, Leibniz-Institute of Vegetable and Ornamental Crops (IGZ), 14979 Großbeeren, Germany; Faculty of Biology & Biotechnology, Ruhr-University of Bochum, 44780 Bochum, Germany

**Keywords:** Xanthomonas, type-III effector, XopM, FFAT-motif, VAP proteins, plant defense

## Abstract

Many Gram-negative pathogenic bacteria use type-III effector proteins (T3Es) as essential virulence factors to suppress host immunity and to cause disease. However, in many cases the molecular function of T3Es remains unknown. The plant pathogen *Xanthomonas campestris* pv. *vesicatoria* (*Xcv*) is the causal agent of bacterial spot disease on tomato and pepper plants and is known to translocate around 30 T3Es into its host cell, which collectively suppress plant defense and promote infection. XopM is an *Xcv* core T3E with unknown function that has no similarity to any other known protein. We found that XopM interacts with vesicle-associated membrane protein (VAMP)-associated proteins (VAPs) in an isoform specific manner. The endoplasmic reticulum (ER) integral membrane protein VAP is a common component of membrane contact sites involved in both tethering and lipid transfer by binding directly to proteins containing a FFAT [two phenylalanines (FF) in an acidic tract (AT)] motif. Sequence analyses revealed that XopM displays two FFAT motifs that cooperatively mediated the interaction of XopM with VAP. When expressed in plants, XopM supports growth of a non-pathogenic bacterial strain and dampens the production of reactive oxygen species, indicating its ability to suppress plant immunity. Further analyses revealed that the interaction with VAP and the ability to suppress PTI are structurally and functionally separable. Although XopM requires localization to the host membrane system for full PTI suppression activity. We discuss a working model in which XopM uses FFAT-motifs to target the membrane in order to interfere with early PTI responses.

## 1. Introduction

Plants can detect attempted invasion by microbial pathogens by perceiving microbe-associated molecular patterns (MAMPs) through transmembrane cell surface-localized pattern recognition receptors (PRRs). Upon activation by MAMP binding, PRRs initiate a number of cellular responses, including the generation of reactive oxygen species (ROS), the initiation of MAP kinase signaling, production of defense hormones such as salicylic acid (SA), and large-scale transcriptional reprogramming followed by the production of antimicrobial proteins and metabolites, as well as cell wall fortification by callose deposition at the cell - pathogen interface (DeFalco & Zipfel, 2021). Collectively, these responses are sufficient to prevent multiplication and spread of a broad range of potential pathogens and thus result in MTI (MAMP-triggered immunity). Plant bacterial pathogens mostly multiply within the apoplastic space of the infected tissue and therefore an effective defense response is heavily dependent on the extracellular release of defense-related molecules, such as pathogenesis-related proteins, secondary metabolites and cell wall materials. Thus, it does not come as a surprise that secretory and endocytic membrane trafficking pathways lie at the heart of plant’s immune system (Jeon & Segonzac, 2023, Gu *et al*., 2017, Yun & Kwon, 2017).

The plant early secretory pathway comprises the ER and the numerous cisternal stacks of the Golgi apparatus. As such, the ER plays a pivotal role in plant immunity by serving as a key site for the synthesis, folding, and quality control of proteins involved in defense responses, thus ensuring the proper functioning of immune receptors and defense molecules (Schäfer & Eichmann, 2012). The ER establishes direct membrane contact sites with the plasma membrane at so called ER – plasma membrane contact sites (EPCS). In plants, EPCSs serve as critical hubs for phospholipid homeostasis and cell integrity (Schapire *et al*., 2008, Ruiz-Lopez *et al*., 2021), are involved in endocytosis (Stefano *et al*., 2018) and autophagy (Wang *et al*., 2019), and regulate cell–cell transport at plasmodesmata (Levy *et al*., 2015, Ishikawa *et al*., 2020).

Vesicle-associated membrane protein (VAMP)-associated proteins (VAPs) are type II ER integral proteins which are conserved amongst phylogenetically distinct organisms and work as tethers between the PM and EPCSs (Stefan *et al*., 2011). As type II ER integral proteins VAPs are inserted into the ER membrane via their C-terminus; however, they lack a domain for making contact *in trans* with the opposing membrane. Thus, it is assumed that VAPs rely on the interaction with other proteins to form multi subunit complexes and hence function as hubs to recruit numerous proteins to the ER (Murphy & Levine, 2016). The architecture of VAPs is modular and composed of three conserved domains: an N-terminal major sperm protein (MSP) domain followed by a central amphipathic domain that is predicted to comprise a coiled-coil domain, and a C-terminal hydrophobic transmembrane domain.

One primary class of VAP-binding partners are cytoplasmic proteins containing two phenylalanines (FF) in an acidic tract (the FFAT motif), which binds specifically to the MSP domain (Loewen & Levine, 2005, Kamemura & Chihara, 2019). The canonical FFAT motif has a core of seven residues, E-F-F-D-A-x-E in single letter amino acid code, and an adjacent upstream region with multiple acidic or phosphorylated residues that additionally contribute to the motif (Loewen & Levine, 2005, Murphy & Levine, 2016, Slee & Levine, 2019). However, most verified FFAT motifs deviate from the canonical motif in their core and/or upstream residues, indicating that a certain degree of sequence flexibility does not necessarily affect FFAT motif functionality (Slee & Levine, 2019). In yeast, four VAP interacting proteins have experimentally been shown to contain FFAT motifs and bioinformatics analysis suggest that the yeast proteome might contain up to 55 FFAT containing proteins capable of VAP binding (Slee & Levine, 2019). Although a number of VAP interacting proteins have recently been identified in plants (Codjoe *et al*., 2022, Greer *et al*., 2020, Fox *et al*., 2020, Stefano et al., 2018, Saravanan *et al*., 2009, Wang et al., 2019, Wang *et al*., 2014), the role of FFAT motifs in mediating the binding of plant VAPs to their alleged protein partners is unknown as none of these proteins has yet been reported to contain a functional FFAT motif.

In Arabidopsis, the VAP family comprises 10 members (VAP27-1 to VAP27-10) divided into three distinct phylogenetic clades (Wang *et al*., 2016). Members of clade I and III possess a single TM domain that determines their localization to ER, while clade II members lack this domain and are assumed to localize to the PM (Wang et al., 2016). Clade I VAPs VAP27-1 and VAP27-3 were shown to localize to the ER as well as to EPCSs (Wang et al., 2014, Wang et al., 2016) and demonstrated to bind to clathrin at endocytic membranes (Stefano et al., 2018).

Although VAPs appear central to membrane processes involved in pathogen defense, a general role for VAPs in plant immunity has so far not been established.

In order to overcome MTI and to cause disease on a given host plants, adapted pathogens have evolved virulence factors that act to suppress plant immunity and promote infection. The injection of type III effector (T3E) proteins into the host cell is an efficient mechanism employed by many Gram-negative bacterial pathogens to overcome MTI and establish a favorable environment (Büttner, 2016, Khan *et al*., 2018, Macho & Zipfel, 2015). T3Es encode proteins with diverse biochemical activities which interfere with host cellular processes on all levels of the plants immune response, including ER – mediated processes or vesicle trafficking (Jeon & Segonzac, 2023, Yun *et al*., 2023).

The Gram-negative bacterium *Xanthomonas campestris* pv. *vesicatoria* (*Xcv*; synonymously designated as *Xanthomonas euvesicatoria*) is the causal agent of bacterial spot disease in pepper and tomato plants (Jones *et al*., 1998). The ability of *Xcv* to cause disease is largely dependent on a suite of approximately 35 T3Es several of which are conserved between different *Xcv* strains (Potnis *et al*., 2011, Schwartz *et al*., 2015). The *Xcv* effectors XopB and XopJ localize to host cell membranes and were shown to specifically inhibit MAMP – induced callose deposition and block secretion of a GFP reporter fused to a signal peptide (Bartetzko *et al*., 2009, Schulze *et al*., 2012, Priller *et al*., 2016). The protein XopM has been identified as a type-III secreted effector in *Xcv* (Schulze et al., 2012) and has later been shown to interfere with MAMP – induced reporter gene expression in a protoplast system (Popov *et al*., 2016). XopM does not share significant sequence similarity with other effector proteins or with any protein found in databases and its biochemical activity, target protein in the host cell, and contribution to *Xcv* virulence remains unknown. In this study, we show that XopM interacts with plant VAPs in an isoform specific manner via two FFAT motifs within the N-terminal part of the effector. XopM suppresses PTI responses when ectopically expressed in plants; however, the ability to affect plant defense was shown to be structurally separable and functionally independent from the VAP binding activity. Further protein-protein interaction studies suggest that XopM interacts with key players of membrane trafficking regulation. Thus, we discuss a model in which XopM uses FFAT motifs as targeting signals to reach certain sub-compartments of the host endomembrane system in order to interfere with vesicular transport processes.

## 2. Results

### 2.1. XopM suppresses plant immunity

To test the contribution of XopM to *Xcv* virulence under laboratory conditions, we generated a *ΔxopM* deletion mutant strain in the *Xcv* 85-10 background and subsequently compared bacterial growth relative to the wild type strain in susceptible pepper plants. Six days after inoculation the *ΔxopM* knockout strain reached similar cell titers as the control (Supplementary Figure S1), indicating that the removal of this single effector protein from the overall *Xcv* effector repertoire does not affect bacterial virulence under the chosen experimental conditions.

To test the effect of XopM expression in plant cells, stable transgenic Arabidopsis lines expressing a XopM - green fluorescent protein (GFP) fusion protein under the control of a *β*-estradiol inducible promoter were generated and expression of the fusion protein was verified by immunoblotting (Supplemental Figure S2). Independent transgenic Arabidopsis lines supported enhanced bacterial growth of a nonpathogenic *ΔhrcC P. syringae* DC3000 (*Pst* DC3000) strain, strictly depending on the induction of XopM-GFP expression by β-estradiol treatment (Figure 1). This indicates that XopM-GFP interferes with MTI responses in Arabidopsis.

**FIGURE 1.**
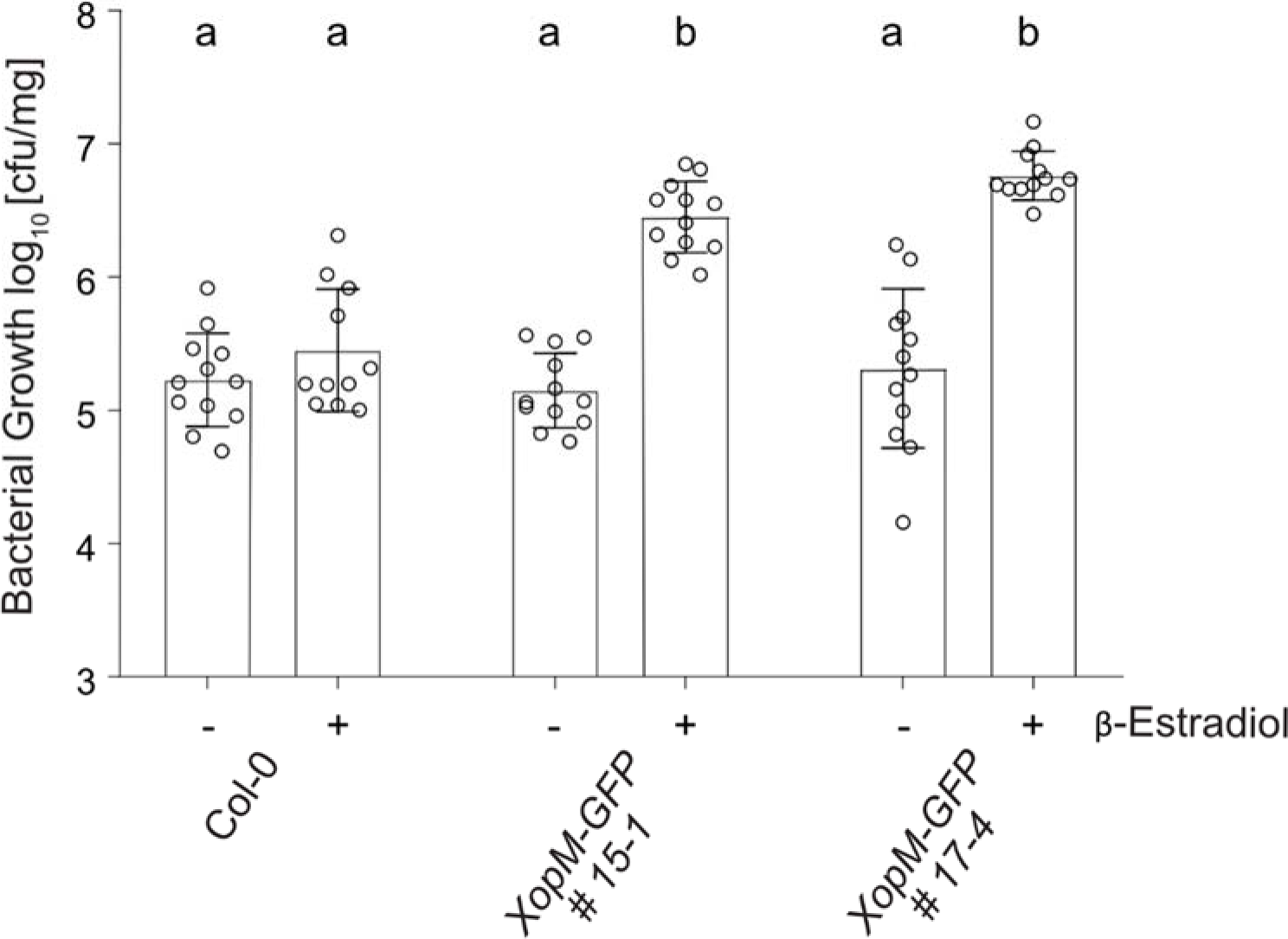
Bacterial growth of XopM-GFP expressing Arabidopsis plants. XopM expression supports growth of the T3SS deficient bacteria *Pst ΔhrcC*. Two-week-old Col-0 and XopM-GFP transgenic Arabidopsis seedlings were either treated with 50 µM β-Estradiol (+) to induce XopM-GFP production or with 0.1% Ethanol (-) as a control. The following day, the seedlings were infected by flooding with *Pst ΔhrcC* (5×10^6^ CFU ml^−1^). The colony forming units per mg of plant tissue (cfu/mg) was measured three days post infection (3 dpi). Individual values are plotted with the bar representing the mean of n ≥ 10. Error bars indicate the standard deviation. Letters indicate statistical significance (p < 0.05) calculated by a one-way ANOVA with a Tukey’s multiple comparison test. The experiment was carried out three times with similar results.

Immune responses generally triggered by MAMPs include a burst of reactive oxygen species (ROS) (Kadota et al., 2015). When transiently expressed in leaves of *Nicotiana benthamiana* using *Agrobacterium tumefaciens* (hereafter, Agrobacterium), XopM fused to GFP at its C-terminus suppressed a ROS burst triggered by flg22 treatment to a similar extent as the positive control AvrPto from *Pst* DC3000 (Figure 2A). In a complementary approach, we tested the ability of XopM to suppress the activity of extracellular peroxidase (POX) enzymes which are induced upon MAMP perception (Mott et al., 2018). Similar to the observation before, XopM was able to suppress POX activity almost down to the level of the negative control (Figure 2B). Expression of all proteins was verified by immunoblotting (Supplementary Figure S3).

**FIGURE 2.**
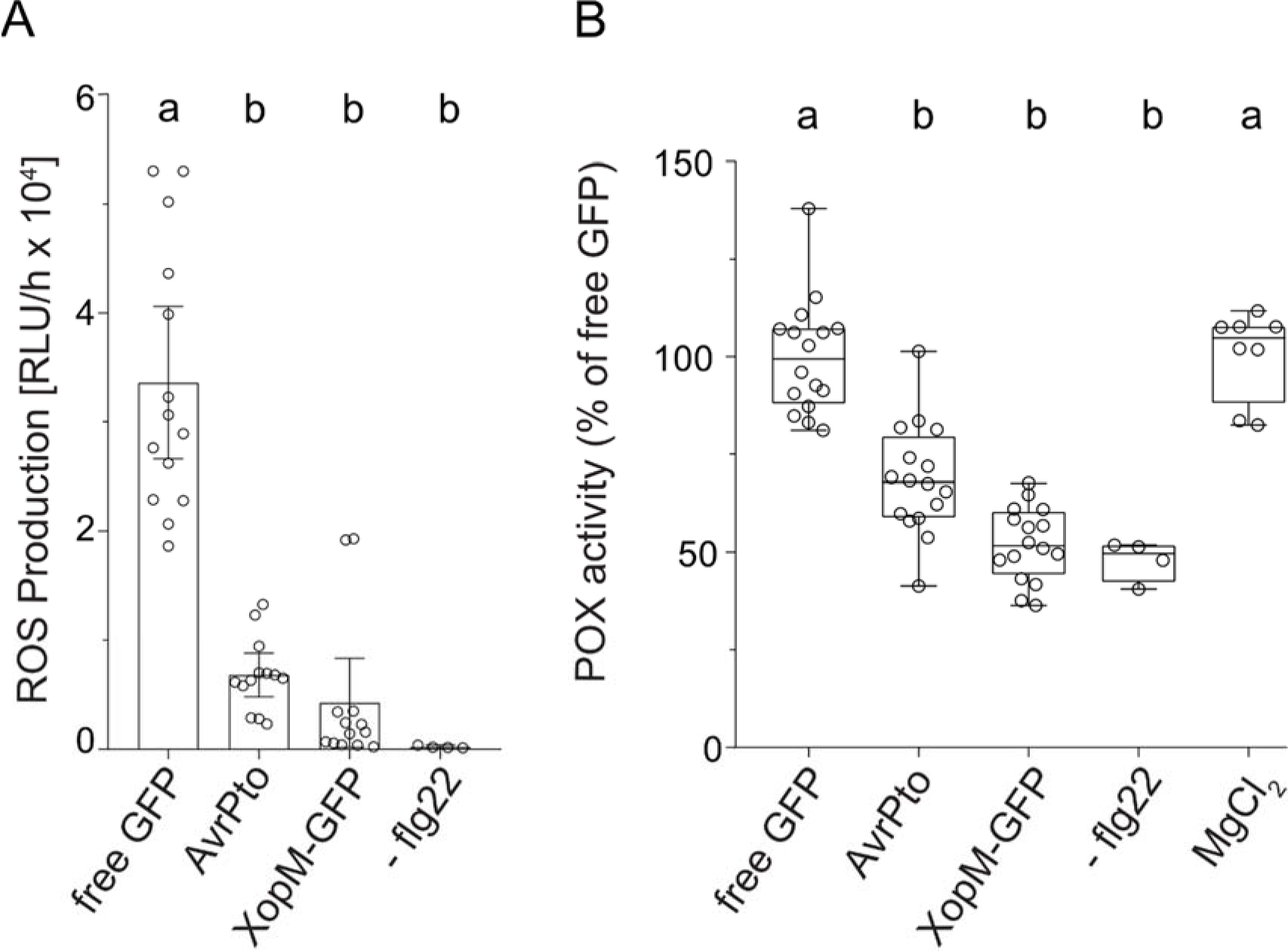
XopM interferes with ROS production and suppresses flg22 induced peroxidase (POX). (A) flg22-induced production of reactive oxygen species (ROS) was measured in leaf discs transiently expressing effector proteins using a luminol based assay. Leaf discs were treated with flg22 and ROS production measured as relative light units (RLU) over the time course of an hour. Leaves infiltrated with MgCl_2_ were not treated with flg22 as a negative control (-flg22). At least twelve samples were measured (n ≥ 12), except for the negative control where only four samples were measured (n = 4). Individual values are plotted with bars representing the mean and error bars give the standard deviation (±SD). Statistical significance was calculated by one-way ANOVA with a Tukey’s multiple comparison test. The different letters indicating significant differences (p < 0.05). (B) Relative peroxidase (POX) activity of flg22 treated leaf discs transiently expressing GFP-tagged effector proteins. Leaf discs were treated with 100 nm flg22 for 16 h and POX activity was measured. Discs infiltrated with MgCl_2_ were not treated with flg22 (-flg22). At least sixteen leaf discs (n ≥ 16) were measured, except for the negative control where n = 8. POX activity of free GFP was set to 100 and the relative activity of the others adjusted to it. Different letters indicate a significant difference (p < 0.05). One-way ANOVA with a Tukey’s multiple comparison test was used. Boxes show the lower quartile, median and upper quartile values. Error bars represent 95 % confidence interval. The experiments were carried out three times with similar results.

Taken together, these findings indicate that despite the fact that the individual deletion of XopM from *Xcv* does not affect bacterial growth under laboratory conditions, XopM possess substantial MTI suppression activity in different plant species. Therefore, it appears highly likely that XopM targets are conserved in different plants irrespective of their susceptibility towards *Xcv*.

### 2.2. XopM interacts with VAP proteins in yeast and in planta

We next aimed to identify the cellular targets of XopM in the plant cell. To this end, we screened a yeast two-hybrid (Y2H) cDNA library from tobacco (*Nicotiana tabacum*) for proteins that interact with XopM. Although tobacco is not a *Xcv* host species, the library has proven very useful to identify *Xcv* T3E target proteins (Üstün *et al*., 2013, Leong *et al*., 2022, Raffeiner *et al*., 2022). A screen of <1 x 10^6^ CFU yielded two candidate XopM interaction partners and a sequence analysis revealed that both encode for VAP proteins. One was identical to VAP12 from *Nicotiana tomentosiformis* (GenBank Acc. XM_009796889.1), the other one matched VAP12 from *Nicotiana sylvestris* (GenBank Acc. XP_009627005.1) when compared to the protein database. Thus, both sequences likely represent homeologous genes derived from the two parental genomes of *N. tabacum*, which is an allotetraploid. A direct interaction assay in yeast between XopM and both VAPs confirmed the interaction (Figure 3A). Both proteins share 97% similarity on the amino acid sequence level with each other and thus we decided to proceed with only one of the clones for further analyses (Acc. XM_009796889.1), which is subsequently named *Nt*VAP12.

**FIGURE 3.**
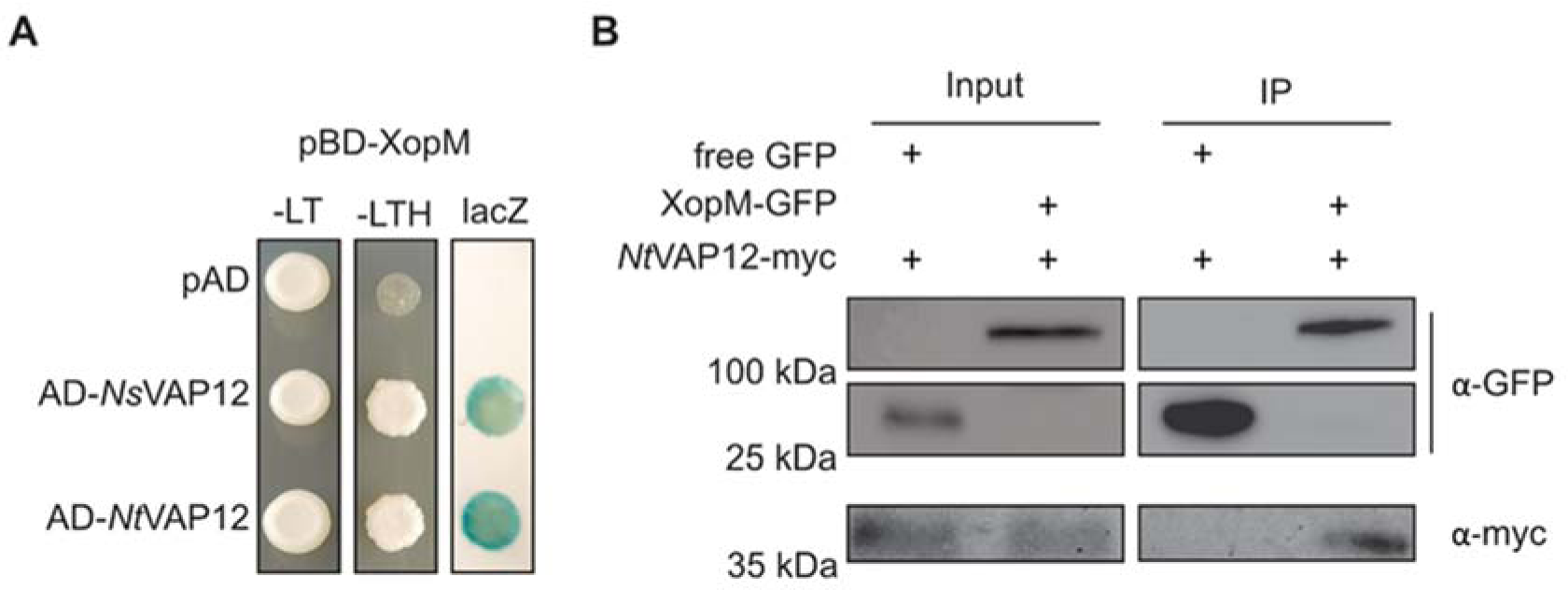
The type-3 effector XopM interacts with VAP12. (A) XopM fused to the GAL4 DNA binding domain (BD) was co-expressed with two tobacco VAP12 proteins fused to the GAL4 activation domain (AD). The empty vector containing the AD (pAD) was used as a negative control. – LT, yeast growth on medium without Leu and Trp. – HLT, yeast growth on medium lacking His, Leu, and Trp, indicating expression of the *HIS3* reporter gene. LacZ, activity of the *lacZ* reporter gene. (B) Confirmation of XopM VAP12 interaction *in planta* via co-immunoprecipitation (co-IP). Using *Agrobacterium*-mediated transient expression, *Nt*VAP12-myc was transiently co-expressed with free GFP or XopM-GFP. After 24 h the total protein content (Input) was treated with GFP-beads (IP). Recombinant proteins were detected by immunoblotting using anti-GFP or anti-myc antibodies. *Ns, N. sylvestris*; *Nt*, *N. tabacum*.

To determine if XopM interacts with *Nt*VAP12 *in planta*, a GFP pull-down assay was performed. To this end, we transiently expressed either XopM-GFP or free GFP each combined with or without *Nt*VAP12-myc *in N. benthamiana*. One day after infiltration with *Agrobacteria*, we performed a pull-down of XopM-GFP using GFP-Trap beads and analyzed the eluates by immunoblot-ting with anti-GFP and anti-myc antibodies. XopM-GFP, but not free GFP, was able to pull down *Nt*VAP12-myc, verifying the specific interaction of both proteins *in planta* (Figure 3B).

### 2.3. XopM and VAP12 colocalize to EPCS

To study XopM protein intracellular localization relative to the localization of *Nt*VAP12, XopM and *Nt*VAP12 were tagged with mCherry and GFP, respectively, and transiently expressed in *N. benthamiana* leaves. XopM-mCherry signals were found close to the PM with very little overlap with the free GFP control (Figure 4A), further substantiating a possible membrane association of XopM. The *Nt*VAP12-GFP was most apparent close to the PM and in a reticular network, characteristic of the ER (Figure 4B). In addition, *Nt*VAP12-GFP fluorescence was also visible in in punctuated structures, as previously described for *At*VAP27s present in EPCS (Wang et al., 2014, Wang et al., 2016). When co-expressed, the XopM-mCherry and the *Nt*VAP12-GFP signals displayed considerable overlap, including the regions representing possible EPCS (Figure 4A).

**FIGURE 4.**
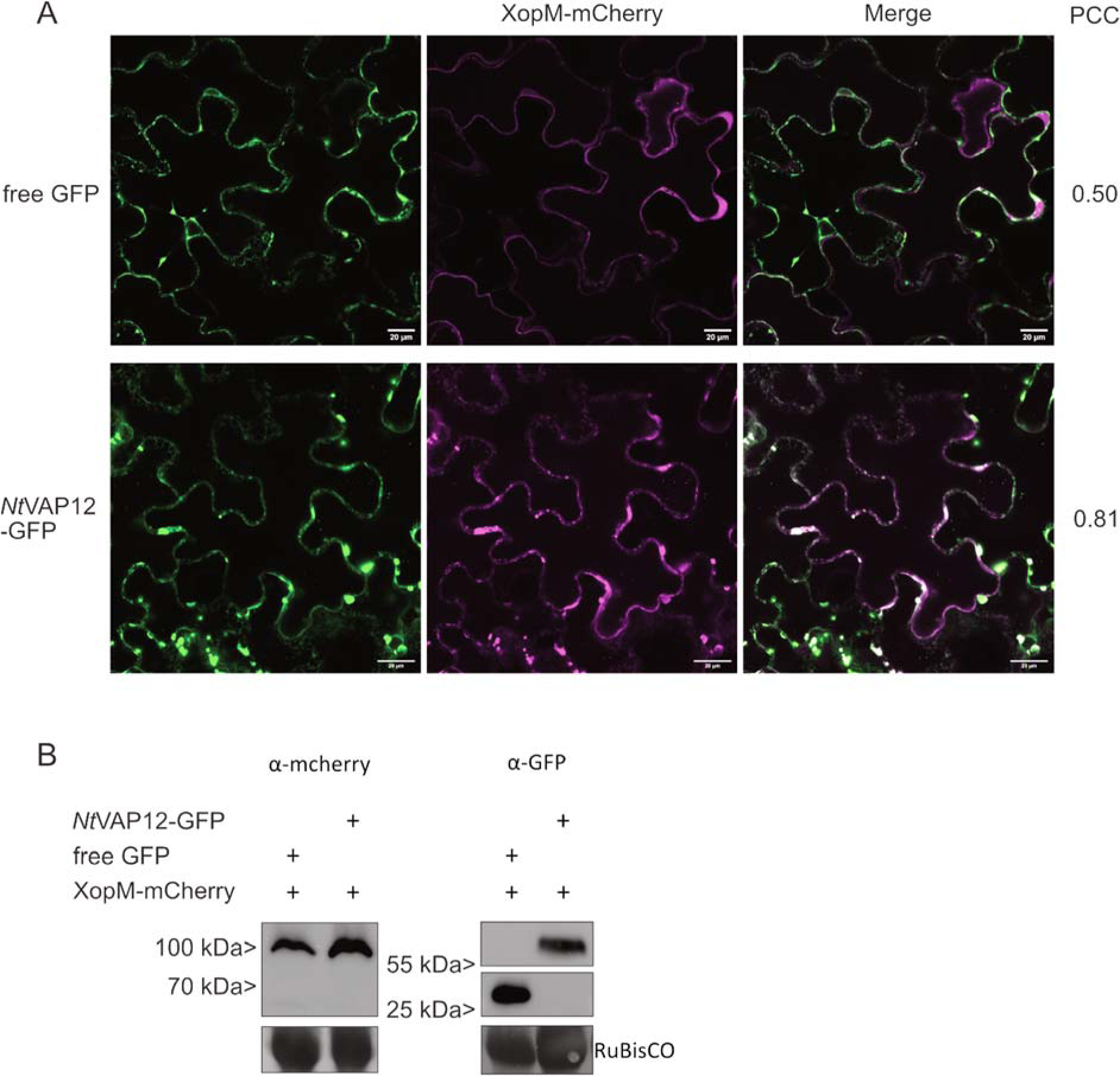
XopM-mCherry co-localises with *Nt*VAP12-GFP in plant cells. (A) XopM-mCherry was transiently co-expressed in *N. benthamiana* with either free GFP or *Nt*VAP12-GFP. The localisation of the proteins was detected 48 h after infiltration using a confocal laser scanning microscope. Co-localisation was quantified using the Pearson’s correlation coefficient (PCC). White: Co-localisation of the expressed proteins. Scale bar represents 20 µM. (B) Western blots of co-localisation. RuBisCO stained with AmidoBlack serves as loading control.

Taken together, both fusion proteins localize to similar compartments of the cell within the endomembrane system, including EPCS. This further supports the notion of an *in vivo* interaction of both proteins in plant cells.

### 2.4. XopM interacts with VAPs from different plant species

The Arabidopsis genome encodes 10 VAP isoforms that phylogenetically fall into three different clades (Wang et al., 2016). Based on sequence similarity, the clade I VAPs *At*VAP27-1 and *At*VAP27-3 are the most closely related orthologs relative to *Nt*VAP12 and both interact with XopM in yeast (Supplementary Figure 4A and B). In addition, the clade III Arabidopsis VAP isoform *At*VAP27-4 also binds to XopM in yeast, while another clade I isoform, *At*VAP27-4, does not interact (Supplementary Figure 4B).

We also tested VAP12 homologs from other species for the ability to bind XopM in yeast. As shown in Figure S4B, *N. benthamiana Nb*VAP12 and *Sl*VAP12 from tomato also interact with XopM. In a similar vein, *N. tabacum Nt*VAP2-1, a clade III member most closely related to *At*VAP27-4 also binds XopM in yeast (Supplementary Figure 4B).

The data suggest that XopM interacts with VAPs from different species from at least two of the three phylogenetic clades.

### 2.5. XopM – VAP interaction depends on the FFAT binding domain of VAP

Structural predictions as well as mutation experiments have shown that the majority of VAP interaction partners bind via conserved residues within the MSP, especially those that carry a FFAT motif (Loewen & Levine, 2005, Slee & Levine, 2019). In order to map NtVAP12 residues critical for its interaction with XopM, a *Nt*VAP12 truncation variant limited to the predicted MSP domain ranging from residue 8 to residue 129 of the protein (MSP*_Nt_*_VAP8-129_; Figure 5A), was tested for binding XopM in yeast. As shown in Figure 5B, the *Nt*VAP12 MSP domain was sufficient for interacting with XopM. Multiple residues within the yeast Scs2 VAP protein, including T41 and T42, critical for binding protein partners especially via their FFAT motif were previously identified (Loewen & Levine, 2005). Based on sequence comparison, the corresponding residues in *Nt*VAP12 are T46 and T47. Alanine substitution of T47 abolished the interaction of *Nt*VAP12 with XopM while the substitution of T46 had no apparent effect on binding between the two proteins in yeast (Figure 5C). Similar to the T47A substitution a T46/47A double exchange also abolished the interaction (Figure 5C).

**FIGURE 5.**
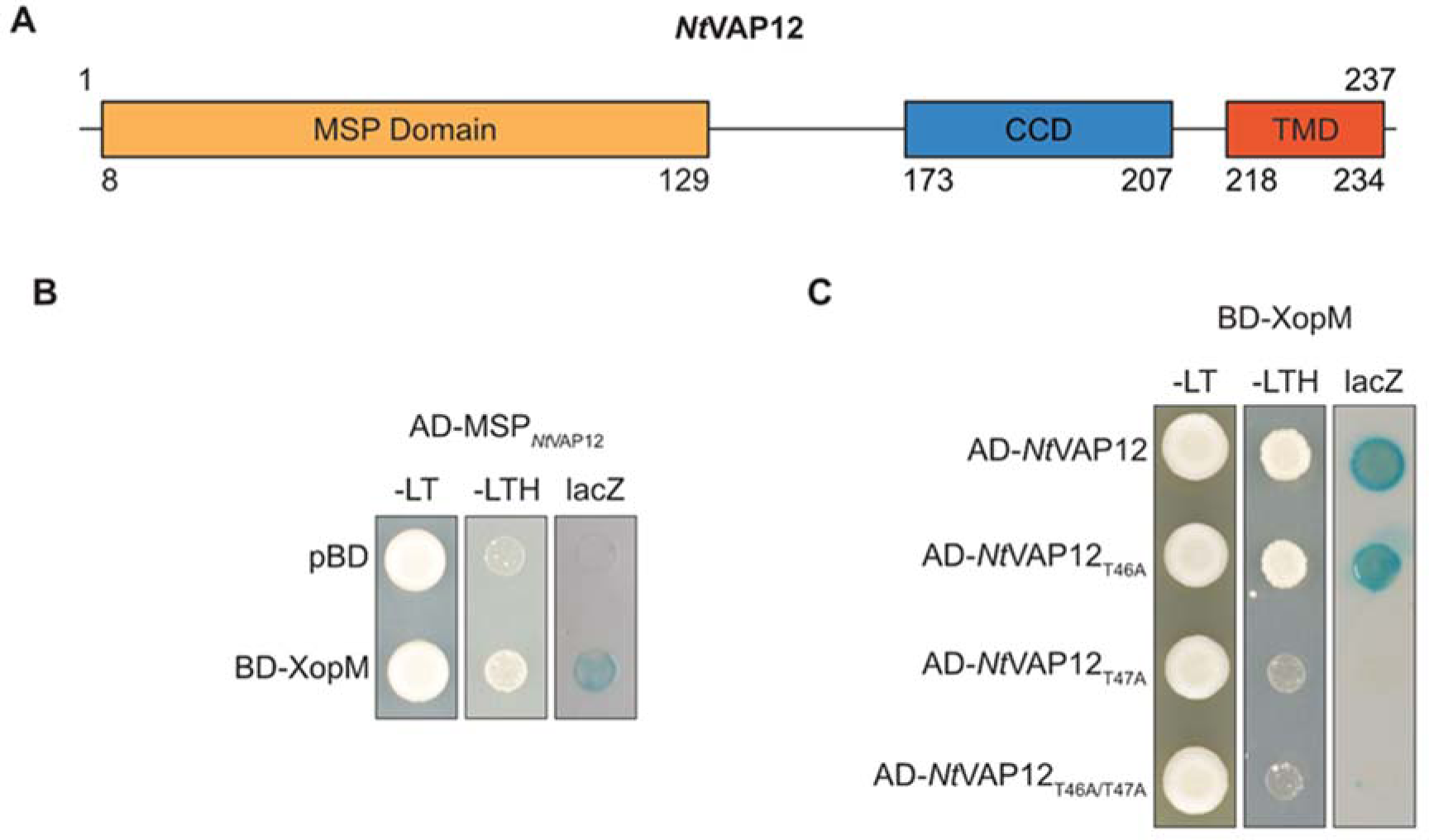
Structure function analysis of the *Nt*VAP12 XopM interaction. (A) Schematic representation of *Nt*VAP12. *Nt*VAP12 has a major sperm (MSP) domain at the N-terminal, in the centre a coiled-coiled domain (CCD) and a transmembrane domain (TMD) at the C-terminal. (B) The MSP domain fused to the GAL4 activation domain (AD) was co-expressed with the GAL4 binding domain (BD) fused to XopM. Empty BD vector (pBD) was used as a negative control. (C) Potential residues for binding XopM of *Nt*VAP12 were substituted and co-expressed with XopM in yeast. Empty AD vector (pAD) was used as a negative control. –LT yeast grown on selective medium lacking Leu and Trp. –LTH, yeast grown on selective medium lacking Leu, Trp and His indicating *HIS3* reporter gene activity. LacZ, activity of the *lacZ* reporter gene. *Nt*, *N. tabacum*

These data suggest that the binding of XopM to *Nt*VAP12 might be mediated by a similar mechanism previously described for the interaction of VAPs with FFAT motif – containing proteins (Slee & Levine, 2019).

### 2.6. XopM contains two FFAT motifs that collectively mediate XopM – VAP interaction

To further explore the structural requirements for the XopM binding to VAPs, an algorithm designed by Murphy and Levine (2016) was used to predict the presence of possible FFAT motifs within the XopM polypeptide sequence. The analysis revealed the presence of two putative FFAT motifs located 23 amino acids apart within the N-terminal region of the XopM protein (Figure 6A). One FFAT motif [G^60^-Y^61^-E^62^-T^63^-A^64^-N^65^-D^66^] is quite degenerated when compared to the consensus FFAT sequence [E^1^(F/Y)^2^(F/Y)^3^D^4^A^5^x^6^E^7^] and can be considered as a non-canonical (nc)FFAT motif, while the other predicted motif [Q^90^-F^91^-Y^92^-D^93^-A^94^-E^95^-D^96^] matches the consensus sequence to a higher degree. Both motifs are preceded by a stretch of serine residues which, if phosphorylated, could provide the acidic environment typically located upstream of the consensus FFAT sequence (Murphy & Levine, 2016). A comparison of non-duplicated XopM sequences lodged with GenBank (185 sequences in total) indicates conservation of both FFAT motifs across *Xanthomonas* species and strains (Figure 6B).

**FIGURE 6:**
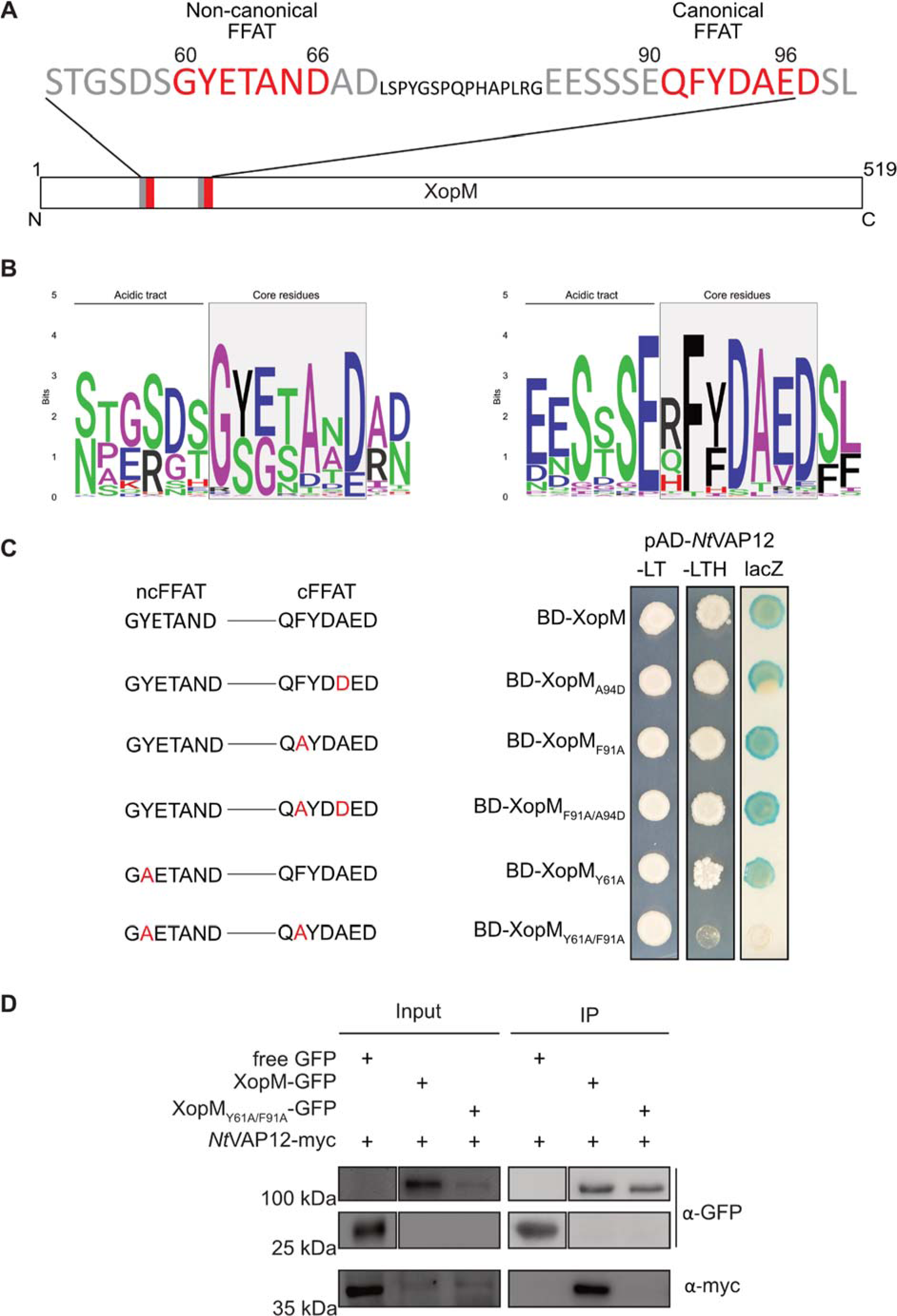
Structure function analysis of XopM. (A) Schematic representation of XopM. XopM has two FFAT (two phenylalanines [<u>FF</u>] in an <u>a</u>cidic tract) at its N-terminal. The non-canonical (ncFFAT) and canonical (cFFAT) motif are highlighted in red, the acidic tract in grey. (B) Representation of the level of conservation of individual residues of the FFAT motifs of XopM. Amino acid properties: green = polar, purple = non-polar, black = aromatic, red = positive, blue = negative. (C) The seven core residues of each motif are shown and the amino acids and the residue that was substituted in each experiment is marked red. The tobacco VAP12 (*Nt*VAP12) fused to the GAL4 DNA binding domain (BD) was co-expressed with different XopM proteins fused to the GAL4 activation domain (AD). –LT yeast grown on selective medium lacking Leu and Trp. –LTH, yeast grown on selective medium lacking Leu, Trp and His indicating *HIS3* reporter gene activity. LacZ, activity of the *lacZ* reporter gene. (D) Confirmation of loss of interaction when mutating FFAT motifs via *in planta* Co-IP. Free GFP, XopM-GFP and XopM_Y61A/F91A_-GFP were co-expressed in *N. benthamiana* with *Nt*VAP12-myc. The total protein content (Input) and after treatment with GFP-beads (IP) were detected by immunoblotting using anti-GFP and anti-myc antibodies. *Nt*VAP12, *N. tabacum*.

Previous data suggest that a phenylalanine or a tyrosine residue at position 2 of the FFAT motif is critical for VAP binding, while amino acid substitutions are tolerated at most other positions (Murphy & Levine, 2016). To test the functional importance of the putative FFAT motifs in XopM, amino acid substitutions were introduced at different positions within the motifs, including the essential position 2 of the ncFFAT (Y61A), the cFFAT (F91A), or both FFAT motifs (Y61A/F91A), as well as a conserved alanine residue at position 5 of the cFFAT (A94D). A direct interaction assay in yeast between *Nt*VAP12 and the respective XopM substitution variant revealed that altering only one of the FFAT motifs does not affect the ability of XopM to bind *Nt*VAP12 (Figure 6C), while the simultaneous change of both FFAT motifs abolishes the interaction. This indicates that at least one intact FFAT motifs is required for the XopM/VAP12 interaction to occur while both motifs appear to be functionally equivalent.

To further corroborate these findings an *in planta* immuno-precipitation experiment was carried out where *Nt*VAP12-myc was co-expressed in combination with either XopM-GFP or XopM_Y61A/F91A_-GFP in leaves of *N. benthamiana*. *Nt*VAP12-myc was pulled down by XopM-GFP only but not by the XopM_Y61A/F91A_-GFP variant (Figure 6D), confirming the functional relevance of the FFAT motifs for XopM binding to *Nt*VAP12 also *in planta*.

### 2.7. Structure prediction of the XopM/VAP protein-protein interaction

AlphaFold, an artificial intelligence (AI) tool created by DeepMind for predicting protein structures, has been recognized as an exceptionally accurate method when compared to other established *in silico* approaches (Fowler & Williamson, 2022). In order to obtain further functional insights into the interaction between XopM and VAP, the AlphaFold Colab was utilized to predict the 3D structure of XopM in isolation as well as complexed with VAP. When considered in isolation, the N-terminal region XopM, harboring the two FFAT motifs, was predicted to be unfolded (Supplementary Figure 5A). Bacterial T3Es typically contain intrinsically disordered regions in the N-terminus that can fold in a stimulus-dependent manner (e.g., during protein–protein interactions) (Marin *et al*., 2013). The remaining part of XopM was predicted to consist of two β sheets and 16 α helices in a bundle – like structure and a comparison to known protein structures using the DALI server (Holm, 2022) did not reveal any further structural clues related to a possible biochemical activity of the effector. However, when XopM was modelled in complex with VAP, the N-terminal region was predicted with high confidence to be associated with the MSP of VAP (Supplementary Figure 5B). Intriguingly, one of the two XopM residues determined to be critical for VAP binding by protein-protein interaction studies (F91) was modelled to be positioned in close proximity to T47 of the MSP of VAP, which in turn is an essential residue for the interaction between the two proteins (Supplementary Figure 5C). These predictions further corroborate a model in which FFAT motifs in XopM mediate the interaction with VAP through conserved T-residues within the MSP.

### 2.8. The N-terminal region of XopM is necessary and sufficient for VAP binding

To further narrow down the structural requirements for binding of XopM to VAP, truncation versions either comprising the XopM N-terminus ranging from residue one to residue 105 and including both predicted FFAT motifs, or encompassing the C-terminal portion of the effector from amino acid 106 to 519 were tested for their ability to interact with *Nt*VAP12 in yeast. As shown in Figure 7A, the N-terminal 105 amino acids of XopM were sufficient to mediate an interaction with *Nt*VAP12 in a Y2H, while the XopM fragment comprising residues 106 to 519 failed to interact with VAP in yeast. This finding was confirmed by an *in planta* pull-down experiment where a myc-tagged *Nt*VAP12 was co-expressed with either XopM_1-106_-GFP or XopM_106-519_-GFP in leaves of *N. benthamiana*. A subsequent immunoblot demonstrated that *Nt*VAP12-myc was precipitated by XopM_1-106_-GFP but not by XopM_106-519_-GFP (Figure 7B).

**FIGURE 7.**
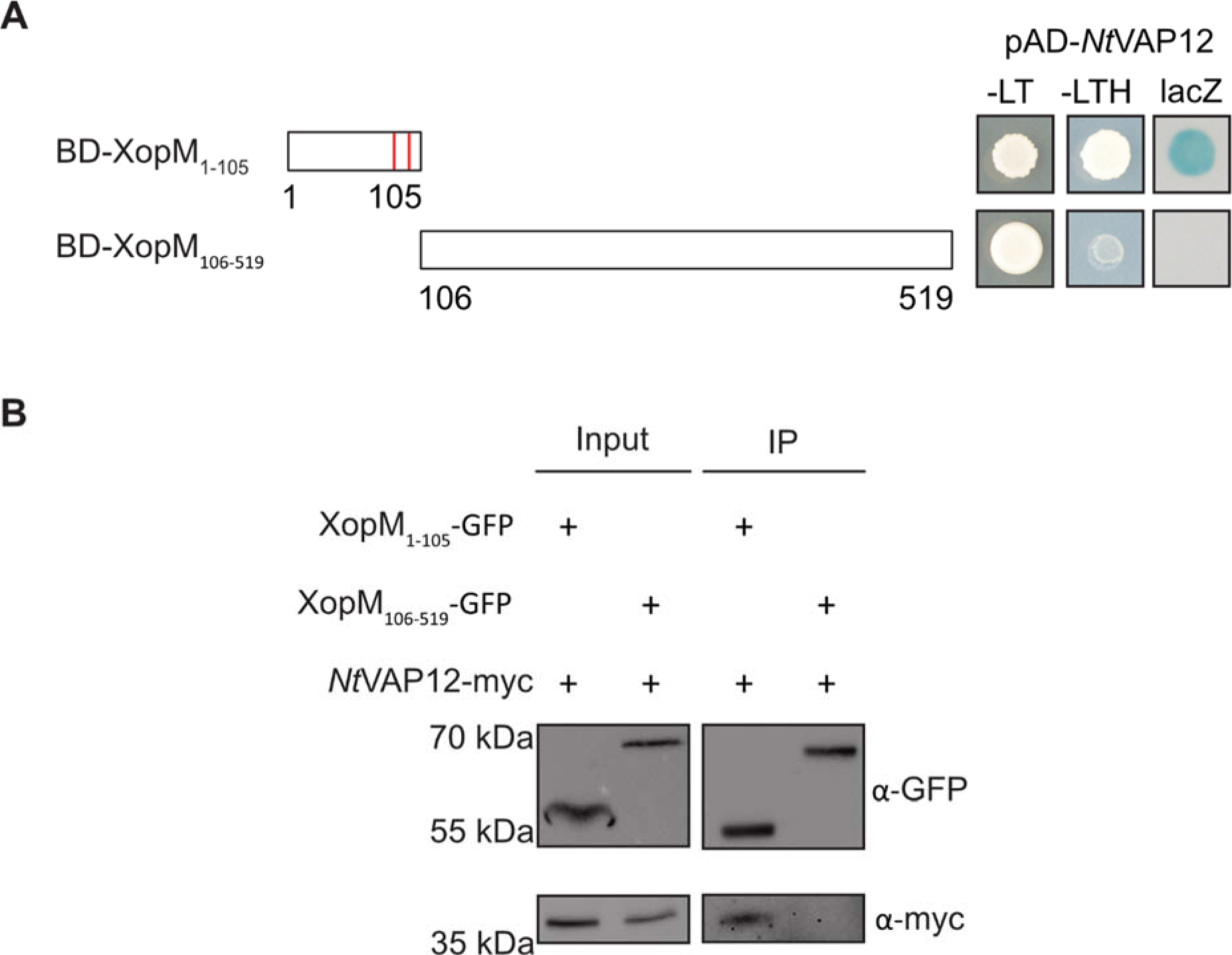
Structure function studies with truncated XopM. (A) Schematic representation of the two truncated variants of XopM: The N-terminal variant (XopM_1-105_) which contains the two FFAT (two phenylalanines [<u>FF</u>] in an <u>a</u>cidic tract) motifs (red dashes) and a C-terminal variant (XopM_106-519_). Proteins were tagged with the GAL4 binding domain (BD) and each co-expressed with *Nt*VAP12 tagged with the GAL4 activation domain (AD). –LT yeast grown on selective medium lacking Leu and Trp. –LTH, yeast grown on selective medium lacking Leu, Trp and His. LacZ, activity of the *lacZ* reporter gene. (B) Co-immunoprecipitation (Co-IP) confirming that only the N-terminal XopM variant that contains the two FFAT-motifs interacts with *Nt*VAP12 *in planta*. XopM_1-105_ and XopM_106-519_ were transiently co-expressed with *Nt*VAP12-myc in *N. benthamiana* leaves. 24h after infiltration the total protein content was extracted (input). After treatment with GFP-beads (IP) detection was carried out by immunoblotting using anti-GFP and anti-myc antibodies. *Nt*VAP12, *N. tabacum*.

These data suggest that the N-terminal region of XopM harboring the two predicted FFAT motifs is necessary and sufficient for binding VAPs.

### 2.9. Virus-induced gene silencing of VAP12 does not affect Xcv growth in *N. benthamiana*

The data obtained so far suggest that XopM specifically interacts with VAP12; however, it is unclear whether this interaction is required to exert its virulence function since a role for VAP12 in plant immunity has far not been established. To further study if VAP12 is involved in the interaction of *Xcv* with its host plant, virus – induced gene silencing (VIGS) with *Tobacco rattle virus* (TRV) of *VAP12* in *N. benthamiana roq1* plants, followed by infection with *Xcv*. ROQ1-deficient plants fail to recognize the *Xcv* T3E XopQ rendering them susceptible to *Xcv* infection (Schultink *et al*., 2017, Gantner *et al*., 2019). For design of the silencing construct, we employed the VIGS tool on the Sol Genomics Network website (Fernandez-Pozo *et al*., 2015) with default settings and using the entire *NbVAP12* coding sequence (CDS) as a query. As a result, the VIGS tool suggested using a fragment covering 300 bp of the *NbVAP12* CDS to achieve high silencing specificity. Strong downregulation of *NbVAP12* of about 90% could be confirmed in *N. benthamiana roq1* plants 2 weeks after infiltration with the VIGS vectors as compared with the pTRV2-*GFP* silencing control (Supplementary Figure 6A). Six plants per construct were selected and used in a bacterial growth assay to investigate if the down regulation of *NbVAP12* effects *Xcv* multiplication. To this end, leaves of pTRV2-*NbVAP12* and pTRV2-*GFP* plants were infected by pressure infiltration with wild type *Xcv* (1×10^5^ CFU mL^−1^) and the bacterial titer was monitored 5 days post infection. As shown in Supplementary Figure S6B, no significant difference in bacterial growth could be observed between *NbVAP12* silenced plants and the *GFP* silencing control. This suggests that *Nb*VAP12 does not play a major role in the interaction between *Xcv* and *N. benthamiana roq1* plants under the experimental conditions applied.

### 2.10. VAP binding and PTI suppression activity of XopM are functionally and structurally separable

The data shown above suggest that the N-terminal 105 amino acids containing both predicted FFAT motifs, are necessary and sufficient for XopM to bind VAP. In order to investigate how the ability of XopM to bind VAP relates to its MTI suppression activity, XopM-GFP truncation variants were transiently expressed leaves of *N. benthamiana*. Measurement of ROS production in response to flg22 revealed that XopM_1-105_-GFP was not able to suppress a flg22 induced ROS burst (Fig. 8A), although the biochemical data suggest that the same N-terminal fragment retained the capacity to bind VAP. In turn, XopM_106-519_-GFP effectively suppressed ROS production, despite loss of VAP binding (Figure 8A). Subcellular localization of XopM_106-519_-GFP suggests that the fusion protein appears still to be associated with the plant cell’s endomembrane system, although the overall localization pattern drastically changes relative to the XopM-GFP wild type protein (Figure 8B). XopM_106-519_-GFP is additionally localized to larger vesicular structures that might resemble endocytotic vesicles. To further delineate whether membrane association per se is required for XopM-GFP to suppress MAMP-induced ROS production, a nuclear localization signal (NLS), forcing the protein into the plant cell nucleus, was introduced into the XopM_106-519_-GFP sequence. Subcellular localization of the NLS-XopM_106-519_-GFP protein confirmed the functionality of the targeting signal *in planta* (Figure 8B). However, the NLS-XopM_106-519_-GFP protein lost the ability to suppress flg22-induced ROS production below the levels of the GFP control (Figure 8A). Expression of all protein variants was confirmed by immuno-blotting (Supplementary Figure 7).

**FIGURE 8.**
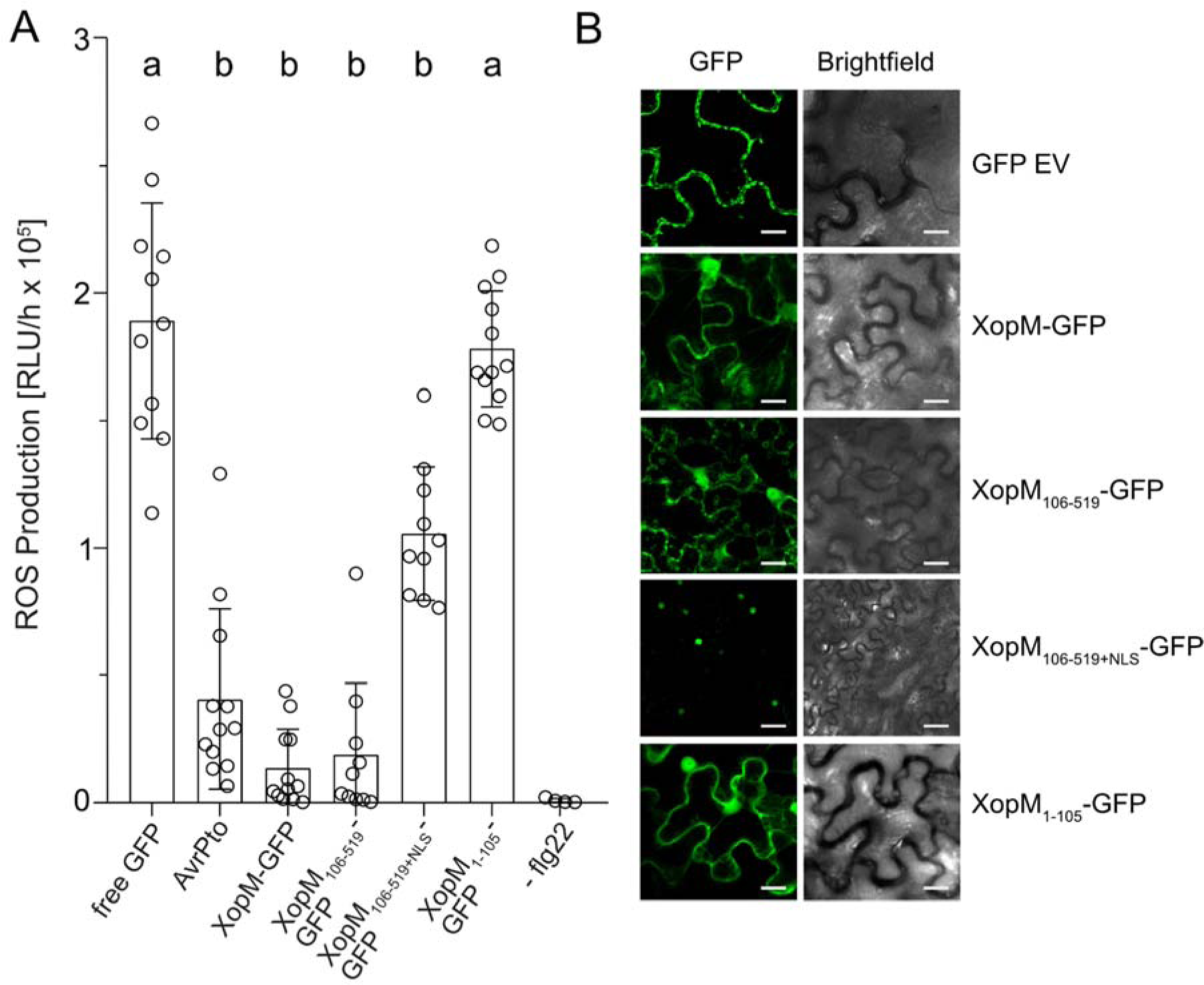
Suppression of flg22 – induced ROS production requires XopM membrane localization but not VAP12 binding. (A) flg22 - induced production of reactive oxygen species (ROS) was measured in leaf discs transiently expressing effector proteins using a luminol based assay. Leaf discs were treated with flg22 and ROS production measured as relative light units (RLU) over the time course of an hour. Leaves infiltrated with MgCl_2_ were not treated with flg22 and served as a negative control (-flg22). At least ten samples were measured (n ≥ 10), except for the negative control where only four samples were measured (n = 4). Individual values are plotted with bars representing the mean and error bars give the standard deviation (±SD). Statistical significance was calculated by one-way ANOVA with a Tukey’s multiple comparison test. The different letters indicating significant differences (p < 0.05). The experiment was repeated three times. (B) Subcellular localization of XopM-GFP variants in *N. benthamiana.* Indicated proteins were transiently expressed in leaves *of N. benthamiana*. The localisation of the proteins was detected 48 h after infiltration using a confocal laser scanning microscope. The scale bar represents 20 µM.

These findings suggest, that at least in a transient CaMV35S-driven overexpression scenario, VAP binding and ROS suppression are functionally independent and structurally separable properties of XopM. However, membrane association appears to be essential for ROS suppression activity of XopM.

### 2.11. Identification of XopM interactors associated with vesicle trafficking

The experiments described above suggest a physical interaction between XopM and VAP in plant cells, however; it appears that XopM virulence function is likely mediated by other cellular interactions. To determine whether XopM has a wider range of interaction partners, we designed a pull-down assay using transient expression of either GFP-tagged *Nt*VAP12, XopM, or XopM_Y61A/F91A_ in leaves of *N. benthamiana*. We queried the interactome of the GFP tagged proteins by co-immunoprecipitation (co-IP) and liquid chromatography-tandem mass spectrometry (LC/MS/MS). Similar to previous reports (Stefano et al., 2018), VAP12 pulled-down cytoskeleton components such as actin and proteins involved in ER membrane trafficking, namely coatomer subunits, dynamin and syntaxin isoforms (Table 1), supporting the validity of the approach. A number of proteins found in the *N. benthamiana* VAPome were annotated to be associated with membrane protein maturation, such as ER membrane protein complex subunits, Sec61 subunits, or calumenin-like proteins. In addition, *Nt*VAP12-GFP associated with other VAP isoforms, indicative for a multimerization of VAPs *in planta*. Of note, the *Nt*VAP12 interactome contained a number of Rab-GTPases (Table 1), which have previously been associated with endocytotic and exocytotic vesicle transport (Nielsen, 2020). A comparison between the VAPome and the XopM interactome revealed a large overlap, although the relative abundance of individual partner proteins varied (Table 1.). For example, the monomeric GTPase RAC5, a known regulator of the cytoskeleton (Klahre *et al*., 2006) was approximately 10 times more abundant in *Nt*VAP12 pull-downs relative to those with XopM or its variant. In turn, VAP isoforms were generally much more abundant in the XopM pull-down relative to the *Nt*VAP12 pull-down, while e.g. calumenin was considerably depleted in the XopM interactome (Table 1.). In contrast to the XopM wild type protein, the XopM_Y61A/F91A_ variant was not able to pull-down VAP isoforms (Table 1), confirming the results obtained from direct interaction assays described above and further supporting the essential role of the XopM FFAT motifs in mediating the XopM/VAP interaction. However, other than the loss of VAP binding, mutation of the two XopM FFAT motifs had a surprisingly little effect on the overall interactome composition when compared to that of the XopM wild type protein. XopM_Y61A/F91A_ still interacted with proteins involved in membrane trafficking, including the group of Rab-GTPases that were found to interact with *Nt*VAP12 as well as with XopM. This indicates that mutation of the XopM FFAT motifs does not qualitatively affect the interaction with other proteins than VAPs and thus might provide an explanation as to why deletion or mutation of the VAP binding region of XopM does not affect XopM’s ability to suppress PTI responses. On a cautionary note, the experimental approach used in the experiments described above involves gross overexpression of XopM in plant cells driven by the CaMV35S promoter, which does not translate well into a natural infection situation where the number of T3E molecules translocated into the host cell is likely less. Thus, overexpression might mask certain effects that are protein level sensitive.

**Table 1.**
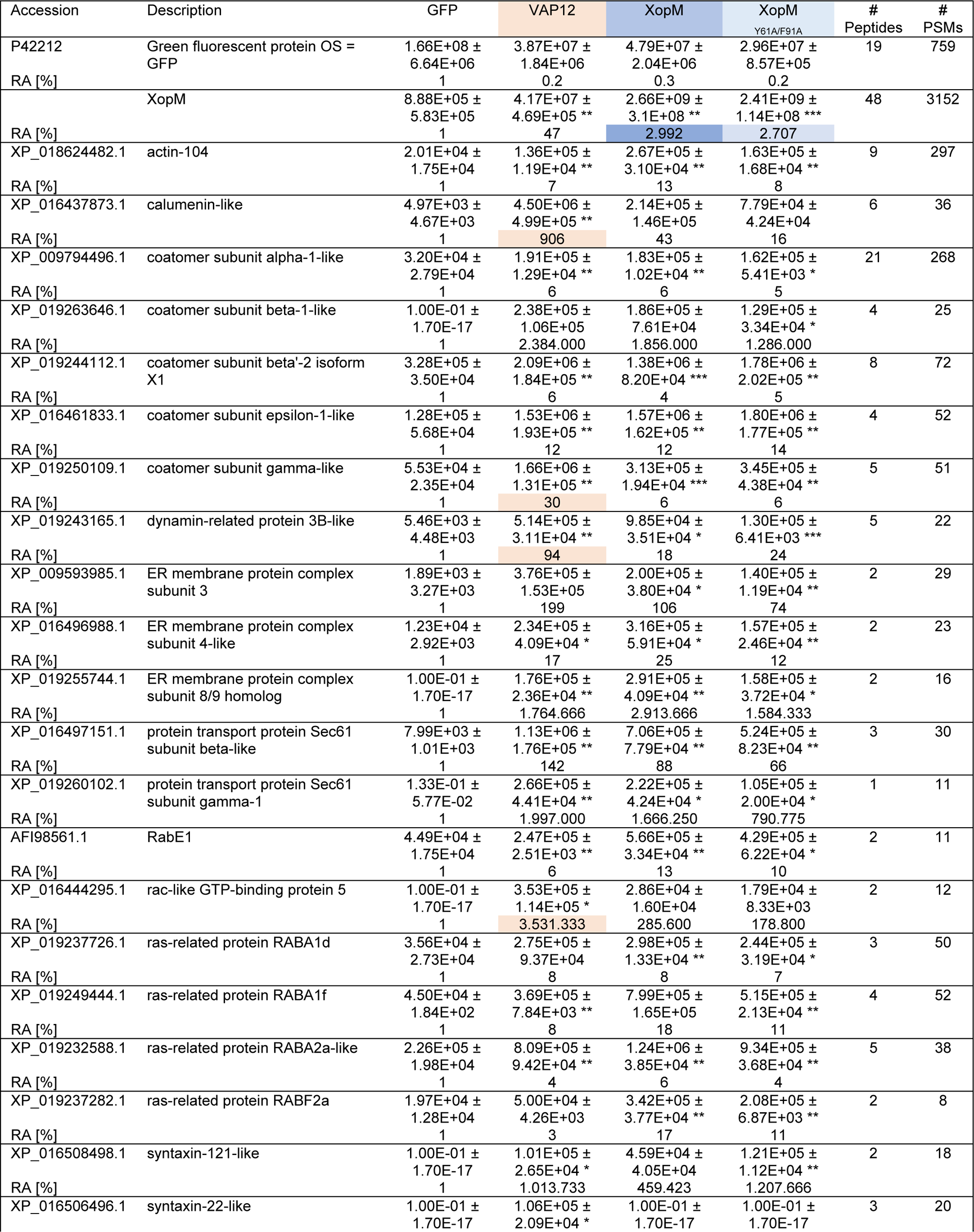
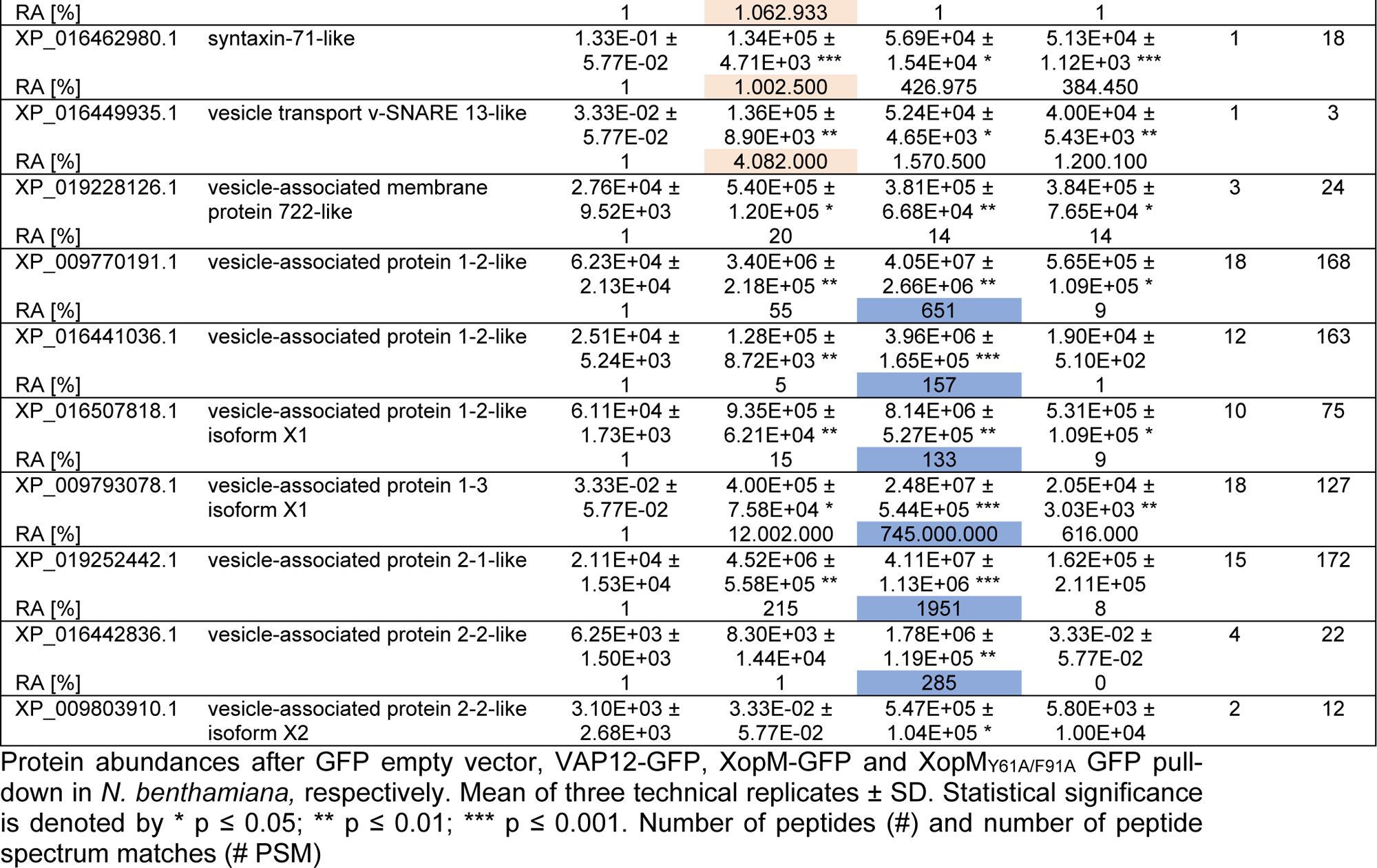
Potential VAP12 and XopM interacting proteins identified by affinity purification coupled to mass spectrometry.

## 3. Discussion

The ability of *Xcv* to cause disease on a given host plant depends on a suite of approximately 35 T3Es, which collectively are essential for bacterial virulence but whose individual contribution to disease progression remains unknown in many cases (Schwartz et al., 2015). Although cellular targets and modes of action are known for only a limited number of *Xcv* effectors, it appears that suppression of MTI responses in the host plant is a major mechanism of effector function (Popov et al., 2016, Büttner, 2016). XopM is a T3E widely distributed among *Xanthomonads* with homologs found in *Pseudomonas* species as well as in *Acidovorax* (Schulze et al., 2012). However, the XopM polypeptide shares no similarity to any other known protein and its function during infection is unknown. An *Xcv xopM* deletion mutant showed no difference in bacterial growth in susceptible pepper plants relative to the *Xcv* wild type strain. It is well known that deletion of individual effector genes often does not result in a change in the virulence phenotype, which may reflect the aspects of the experimental context and/or redundancy of effector function (Castañeda *et al*., 2005). The overlapping activities of bacterial effectors in phytopathogens by targeting the same host targets have been extensively reported (Kvitko *et al*., 2009, Badel *et al*., 2006, Cunnac *et al*., 2011, Munkvold *et al*., 2009).

Therefore, the functional characterization of *Xcv* effectors can benefit from expression of individual effectors either transiently or in stable transgenic plants, rather than studying single effector deletions. Several studies using heterologous expression of individual effectors have revealed their contribution to virulence (Gimenez-Ibanez *et al*., 2018, Hann & Rathjen, 2007, Popov et al., 2016). Inducible expression of XopM in Arabidopsis supports the *in planta* growth of a non-pathogenic *Pst* DC3000 strain, which is normally limited due to MTI. This demonstrates that XopM is able to interfere with MTI responses and can make a significant contribution to bacterial virulence. Bacterial growth assays in XopM expressing plants do not reveal whether a specific MTI response is repressed by the effector; however, transient XopM expression in *N. benthamiana* suggest that the T3E suppresses MAMP-induced ROS production. ROS generation during MTI occurs through PM resident nicotinamide adenine dinucleotide phosphate (NADPH) oxidase family members, which are activated through phosphorylation by multiple protein kinases, including BIK1 (Kadota *et al*., 2014). Upon flg22 recognition, the activated FLS2 complex phosphorylates and releases BIK providing a direct link between MAMP-recognition and elicitation of a ROS burst. Thus, it is conceivable that XopM directly target ROS NADPH-oxidase family members or their direct activation pathway to specifically dampen this branch of the immune response. However, given the fact that XopM also inhibits induction of POX activity points more into the direction of a broader effect of the effector on MTI that could also involve inhibition of defense gene induction or secretion of antimicrobial molecules. This is in line with the observation that XopM inhibits the induction of a flg22 induced defense reporter gene in an Arabidopsis protoplast system (Popov et al., 2016). Several bacterial T3Es have been shown to inhibit PRR activation or downstream signaling and thus impacting multiple immune responses, including ROS production and transcriptional activation (Büttner, 2016). Thus, the inhibition of different defense outputs by XopM might point towards targeting an early step in the signaling cascade, e.g. PRR signaling, at the PM. An at least partial localization of XopM at the PM and/or the endomembrane system is in good agreement with the subcellular localization pattern of the XopM-GFP fusion protein. Several effectors contain consensus sites for myristoylation or palmitoylation at their N-terminus and fatty acid modification has been shown to essential for their localization to the host cell membrane system as well as for their respective virulence function (Bartetzko et al., 2009, Lewis *et al*., 2008, Nimchuk *et al*., 2000). Apparently, XopM does not carry a consensus sequence for fatty acid modification and thus its membrane association must be achieved through different means.

Here, we report the binding of XopM to the ER-membrane tethering protein VAP12 and both proteins display an overlapping localization pattern when expressed in plant cells. VAP proteins are ER-resident tail-anchored membrane proteins and the Arabidopsis orthologs of *Nt*VAP12, VAP27-1 and VAP27-3, have been shown to form ECPS through the interaction with PM localized proteins (Wang et al., 2014, Saravanan et al., 2009). The data shown here suggest that the localization pattern of XopM and *Nt*VAP12 overlaps within ECPS and thus the effector might target a process specific to this particular membrane sub-compartment.

The XopM interaction is specific for certain VAP isoforms and requires the VAP MSD, whereas the CCD and TMD are appear to be dispensable for XopM binding. Mutational analysis within the VAP MSD domain suggest that the XopM interaction depends on residues which have previously been shown to be involved in binding proteins containing FFAT motifs (Loewen & Levine, 2005). Further experiments confirm that XopM contains two eukaryotic FFAT motifs within its N-terminus which are functionally required for the interaction of XopM with VAPs. Intriguingly, both XopM FFAT motifs can substitute for each other for VAP binding suggesting that the presence of two FFAT motifs in XopM is an evolutionary adaptation to increase the interaction with VAPs or represents a possibility to interact with two VAPs simultaneously. The sequence of the XopM cFFAT (Q-F-Y-D-E-A-D) closely matches the consensus FFAT motif and key residues that have been shown to mediate the FFAT/VAP interaction, such as the aromatic amino acid residue at position 2, are conserved (Loewen & Levine, 2005, Furuita *et al*., 2021). Structural predictions of the XopM/VAP interaction suggest that the phenylalanine at position 2 of the XopM cFFAT is positioned in close proximity of critical FFAT interaction residues with the VAP MSP. The ncFFAT of XopM (G-Y-E-T-A-N-S) is more divergent from the FFAT consensus sequence. However, it also carries an aromatic residue at position 2 (Y) and position 4 harbors a threonine residue that could be phosphorylated and therefore mimic the aspartate residue present in the canonical sequence (Di Mattia *et al*., 2020). Furthermore, both XopM FFAT motifs are directly preceded by multiple serine residues, which, if phosphorylated could impart additional negative charges to the motif, which would increase VAP binding (Furuita et al., 2021).

The presence of functional FFAT motifs in XopM is intriguing for several reasons. First, FFAT motifs are considered as eukaryotic motifs generally not present in prokaryotes (Slee & Levine, 2019) and second, functional FFAT motifs in VAP binding proteins from plants have, to our knowledge, so far not been described. Recent work led to the identification of the *Chlamydia trachomatis* Inc protein IncV, which constitutes a structural component that tethers the ER membrane to the membrane of the bacteria containing vacuole inside the host cell through interaction with VAP (Stanhope *et al*., 2017). The IncV–VAP interaction relies on the presence of two FFAT motifs in the C-terminal cytosolic tail of IncV. In analogy to XopM, Incv contains a cFFAT as well as ncFFAT motif with multiple preceding serine residues (Stanhope et al., 2017). Thus, pathogenic bacteria with fundamentally different life styles and virulence strategies appear to employ similar molecular mechanisms to achieve VAP interaction and FFAT motifs in XopM add to the growing arsenal of eukaryotic-like functions in T3Es from plant pathogens as mechanism to mimic host proteins and exert control over host cellular pathways (Büttner, 2016).

Effectors generally operate by binding or enzymatically modifying host molecules. These host molecules are classified into two major functional classes: (1) “targets” directly modified by effectors to modulate host processes; (2) “helpers” or “facilitators” coopted by effectors to execute their activities, for instance to traffic within host cells or to facilitate interaction with target proteins (Win *et al*., 2012). Binding of XopM to VAP is supported by several lines of experimental evidence; however, VAPs do not appear to represent the actual virulence target of the effector. Firstly, VIGS of XopM binding VAPs in *N. benthamiana* has no effect on infection with virulent bacteria and thus, there is currently no substantial evidence that XopM interacting VAPs play an active role in immunity. Secondly, and more importantly, the abilities of XopM to either bind VAP or to suppress MTI responses appear to be functionally independent and structurally separable. While the N-terminal part of XopM, harboring the two FFAT motifs, by itself is able to bind VAP, it has no effect on flg22 induced ROS production when transiently expressed in *N. benthamiana*. In turn, the C-terminal portion of XopM fails to interact with VAP but retains its ability to suppress ROS in an MTI assay. The N-terminally truncated XopM variant still associates with the endomembrane system although the overall localization pattern deviates from that of the full length protein. Only when the T3E is forced into the plant cell’s nucleus MTI suppression is lost. Thus, XopM likely acts at the membrane of the host cell but in at least in a transient overexpression scenario virulence activity does not require VAP binding.

Thus, the question remains what the actual virulence target of XopM might be? A group of proteins that binds to *Nt*VAP12 but also to XopM irrespective of the presence of FFAT motifs, are different isoforms of RAB GTPases. These regulate intracellular transport and the trafficking of proteins between different organelles via endocytotic and secretory pathways (Nielsen, 2020). Several members of this family have been identified to regulated crucial processes of plant immunity. In Arabidopsis, expression of dominant-negative mutants of a number of RABA members affected different steps of endocytosis of FLS2 and thus interfered with MTI activation (Choi *et al*., 2013). Furthermore, the highly conserved effector protein RxLR24, secreted by the oomycete *Phytophthora brassicae* interacts with the Raba GTPase family member RABA1a in vesicles and plasma membranes, inhibiting the function of RABA GTPase in vesicle secretion and reducing vesicle secretion of antimicrobial agents (Tomczynska *et al*., 2018). Thus, it is conceivable and in line with the observations presented here that XopM could target Rab GTPases to interfere with immunity related endocytic or exocytic vesicle trafficking. We currently have no experimental data on a possible biochemical activity of XopM. In principle, XopM binding could interfere with RAB regulation, with the association to the cytoskeleton or with the binding to specific proteins located in target membranes. Future experiments will have to address the functional relevance of the XopM/RAB interaction for the virulence function of the effector.

Intriguingly, the XopM FFAT motif defective mutant still binds to RAB GTPases in an unbiased pull-down experiment while the ability to interact with VAPs is completely lost. This is in line with the observation that loss of VAP binding does not affect XopM virulence function under the experimental conditions but question the relevance of VAP binding of the effector. Given the fact that FFAT motifs are widely conserved in XopM homologs from different Xanthomonads argues for considerable evolutionary pressure for their preservation. Possibly, confined localization of VAP12 in certain membrane sub-compartments, such as EPCS, enables XopM to target specific sub-populations of RAB GTPases with immunity related functions. It has to be borne in mind that ectopic overexpression of effector proteins, as in our study, can led to misleading results by causing promiscuous interactions and regulation of pathways that are associated with the degree of overexpression rather than the function of the protein. Thus, future experiments will have to aim at the functional characterization of XopM protein-protein interactions under more natural conditions. Nevertheless, our study provides novel insights into T3E membrane targeting mechanisms and it will be interesting to determine whether FFAT motifs serve as targeting signals in other *Xanthomonas* T3Es and in effector proteins from other phytopathogens and how this relates to their respective virulence function.

## 4. EXPERIMENTAL PROCEDURES

### 4.1. Plant materials and growth conditions

Pepper (*Capsicum annuum* cv. Early Cal Wonder (ECW)) and *Nicotiana benthamiana* and *N. benthamiana roq1* plants were grown in soil in a growth chamber with daily watering, and subjected to a 16 h light : 8 h dark cycle (25°C : 20°C) at 240-300 µmol m^−2^ s^−1^ light and 75% relative humidity. Transgenic β-estradiol (Sigma-Aldrich) inducible XopM-GFP Arabidopsis (*Arabidopsis thaliana*) lines were generated by transformation of Col-0 wild-type plants with a plasmid containing the entire XopM coding region inserted into binary vector pABindGFP (Bleckmann et al., 2010) using the floral dip method (Clough & Bent, 1998). For the selection of transgenic plants, seeds of T0 plants were sterilized and sown onto Murashige and Skoog medium (Sigma-Aldrich) supplemented with 20 µg/mL hygromycin B (Roth). Primary transformants were allowed to self-fertilize and were propagated into the homozygous T3 generation. Arabidopsis seedlings were grown on ½ MS in an 10h light: 14h darkness (22 °C/18 °C) at a light intensity of 150 μE m^−2^ s^−1^ and 70% humidity. Transgene expression was induced 2 weeks after germination by flooding the petri dishes with 50 µM β-estradiol and 0.1% (v/v) Tween-20. Control plants were flooded with 0.1% (v/v) EtOH in water and 0.1% (v/v) Tween-20.

### 4.2. Bacterial strains and plant infection assays

Strains used in this study were as follows: *Xanthomonas campestris* pv. *vesicatoria 85-10* (*Xcv* wild type), *XcvΔxopM deletion strain, Pseudomonas syringae* pv. *tomato* strain DC3000 *PstΔhrcC* (type-3 secretion-deficient mutant). The *XcvΔxopM* deletion strain was constructed as described (Üstün *et al*., 2013) using the primers listed in Supplementary Table S2. Bacterial infection assays of pepper and *N. benthamiana roq1* plants were carried out exactly as detailed in Raffeiner et al. (2022). The bacterial infection assay of Arabidopsis seedlings was adapted from Ishiga et al. (2011) with a bacterial density of 5×10^6^ CFU ml^−1^. At least three biological replicates were used per experiment unless otherwise stated.

### 4.3. Transient expression of proteins in *N. benthamiana*

Transient expression of proteins using infiltration of leaves of *N. benthamiana* was carried out as described (Raffeiner et al., 2022).

### 4.4. Measurement of ROS production and POX activity

ROS production upon flg22 treatment was monitored using the luminol-based quantification (Liang *et al*., 2013). Measurement of plant peroxidase (POX) was carried out as described (Mott *et al*., 2018) with slight adjustments using a plate reader (Infinite® 200 PRO, Tecan).

### 4.5. Yeast two-hybrid assays

Yeast two-hybrid techniques were performed according to the yeast protocols handbook and the Matchmaker GAL4 Two-hybrid System 3 manual (both Clontech, Heidelberg, Germany) using the yeast reporter strains AH109 and Y187. To identify XopM interacting proteins, a GAL4 AD-domain tagged cDNA library from *Nicotiana tabacum* (Börnke, 2005) was screened as detailed before (Üstün et al., 2013).

### 4.6. *In planta* GFP pull-down assay

Immunoprecipitation was performed by adding 1 g of homogenized leaf material 50 µL of GFP-Trap^®^ coupled to magnetic beads (ChromoTek). Samples were processed as described (Raffeiner et al., 2022).

### 4.7. Immunoblotting

Protein extracts were boiled with 5 x SDS sample buffer and separated by SDS-page. Immunoblotting was carried out with anti-myc antibody (1:2.500, Abcam), followed by Goat Anti-Rabbit IgA alpha chain (HRP) (1:5.000) secondary antibody (Abcam). HA-tagged proteins were detected by anti-HA Peroxidase high affinity antibody (Sigma). GFP was detected using a horseradish peroxidase-conjugated anti-GFP antibody (1:1.000; Santa Cruz Biotechnology Inc., cat. no. sc-9996 HRP). Signals were visualized using chemiluminescence (Thermo Fisher Scientific) with a ChemiDoc Imaging system (Biorad).

### 4.8. Confocal microscopy and image analysis

Confocal microscopy was carried out as previously described (Bortlik et al., 2023). Fluorescence in leaf epidermal cells of *N. benthamiana* was detected by CLSM (LSM880, Axio Observer; Zeiss) after 48 hpi. The specimens were examined using the LD LCI Plan-Apochromat 25x/0.8 water-immersion Imm Korr DIC M27 objective. Fluorescent-tagged proteins were excited by the respective lasers: GFP (ex. 488 nm, em. 507 nm) and mCherry (ex. 587 nm, em. 610 nm). Microscopy images were analysed using Fiji (Schindelin *et al*., 2012). Co-localization was quantified using the JACoP plugin (Bolte & Cordelières, 2006).

### 4.9. Virus – induced gene silencing

VIGS was performed as described previously (Üstün et al., 2013). The mRNA levels of silenced plants were checked via quantitative real-time PCR as described in Raffeiner et al. (2022) using the primers listed in Supplementary Table S1.

### 4.10. Large-scale GFP pull-down and LC-MS/MS analysis

Pull-down followed by LC-MS/MS analysis was basically carried out as previously described (Bortlik *et al*., 2023). In short, on bead trypsin digest of GFP-tagged proteins were treated previously with 100 µl of 0.1 % RapiGest SF (Waters, Eschborn) followed by reduction and alkylation. Trypsin (1:50 (w/w) trypsin/protein) was added and incubated at 37 °C overnight. Subsequently, desalting of peptides was carried out according to Witzel *et al*. (2019). Protein digest were analysed using the Thermo Fisher Q Exactive high field mass spectrometer by reverse-phase HPLC-ESI-MS/MS using the Dionex UltiMate 3000 RSLC nano System coupled to the Q Exactive High Field (HF) Hybrid Quadrupole Orbitrap MS (Thermo Fisher Scientific) and a Nano-electrospray Flex ion source (Thermo Fisher Scientific) as described by Bortlik et al. (2023) with slight modifications. In short, samples were separated on a 25 cm EASY-Spray column (75 µm inner diameter) at 40 °C. Peptides were separated over a linear 100 min gradient of 5-44 % solvent B at a flow rate of 300 nl/min. To identify proteins the mass spectrometer operated from m/z 375-1500 at a resolution of 140 000. Automated gain control (AGC) target was 3 x 10^6^ with a max. injection time of 65 ms. Triplicates of each sample were measured. Scientific) as described previously by Witzel *et al*. (2019). Each sample was measured in triplicate. Proteome discoverer software (PD2.4) was used to analyse and align the LC-MS raw data files, with its built-in MS Amanda, MS Mascot and Sequest HT search engine (Thermo Scientific) (Witzel et al., 2019). The MS data was searched against common contaminants, separated FASTA files of the tagged bait proteins as well as a *N. benthamiana* database based on the Niben1.0.1 draft genome (Kourelis *et al*., 2019).

## Supporting information

Supplementary Figure 1: Deletion of XopM does not affect the virulence of Xcv in pepper (*Capsicum annuum*) plants.

Supplementary Figure 2: Immunoblot analysis of inducible of XopM-GFP expression in transgenic Arabidopsis lines.

Supplementary Figure 3. Verification of effector protein expression for ROS and POX assays.

Supplementary Figure 4: XopM interacts with several VAP isoforms.

Supplementary Figure 5. Structural modelling of XopM and XopM with NtVAP12.

Supplementary Figure 6: Bacterial growth of *Xcv* is not affected by reduced expression of *NbVAP12*.

Supplementary Figure 7. Verification of protein expression of XopM protein variants for ROS assays.

Supplementary Table 1. Oligonucleotides used in this study.

## Acknowledgements

This study was supported by funds from the Deutsche Forschungsgemeinschaft (DFG; German Research Foundation) to F.B. (BO1916/5-2 and CRC973 project C6) and the European Cooperation in Science and Technology EuroXanth (CA16107) from the European Union. S.Ü. was supported by an Emmy Noether Fellowship GZ: UE188/2-1 awarded by the DFG. We thank Kerstin Bieler and Mandy Heinze for their skillful technical help.

## Data availability statement

Any additional data associated to this study are available from the corresponding author upon reasonable request.

## Notes

### Competing Interest Statement

The authors have declared no competing interest.

