## Supplemental Data for "XopM, a FFAT motif containing type-III effector protein from *Xanthomonas*, suppresses PTI responses at the plant plasma membrane"

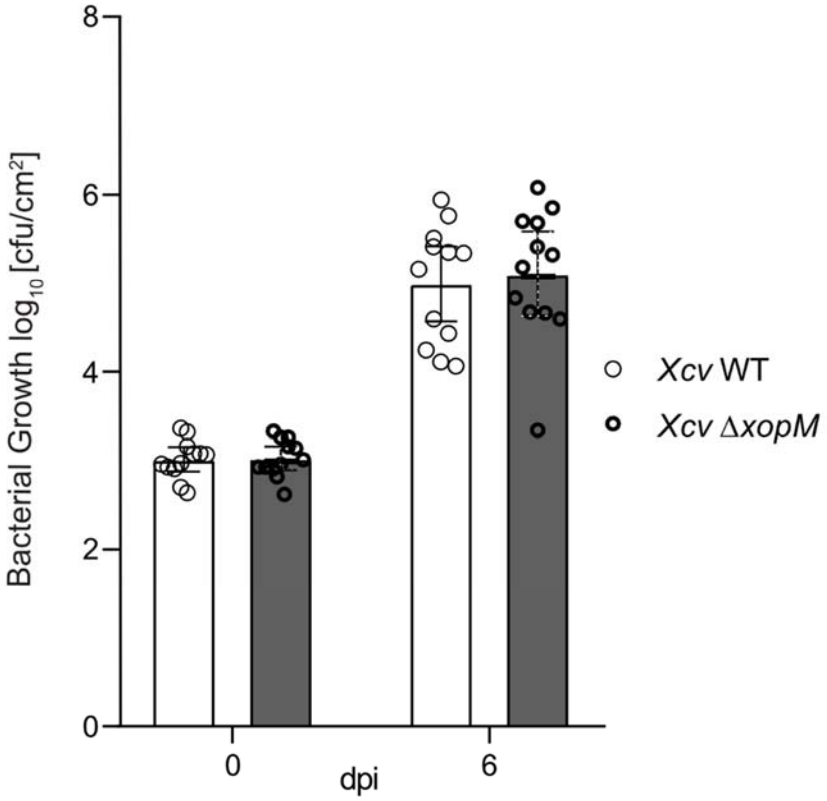


**Supplementary Figure 1: Deletion of XopM does not affect the virulence of *Xcv* in pepper (*Capsicum annuum*) plants.** Leaves of pepper plants (variety ECW) were infected with wildtype (WT) *Xcv* and the XopM knockout strain *Xcv* Δ*xopM*. Leaves were hand-infiltrated with a $10^5$ cells/ml suspension of bacteria. The colony forming units (cfu) were determined on the day of infection (0 dpi) and six days post infection (6 dpi). Individual values are plotted with the bars giving the mean of six biological replicates (and two technical replicates per biological). Error bars indicate the standard deviation (±SD). Statistical difference on each dpi was calculated using a student's t-test with no significant difference ($p>0.05$). The experiment was carried out at least three times with similar results.

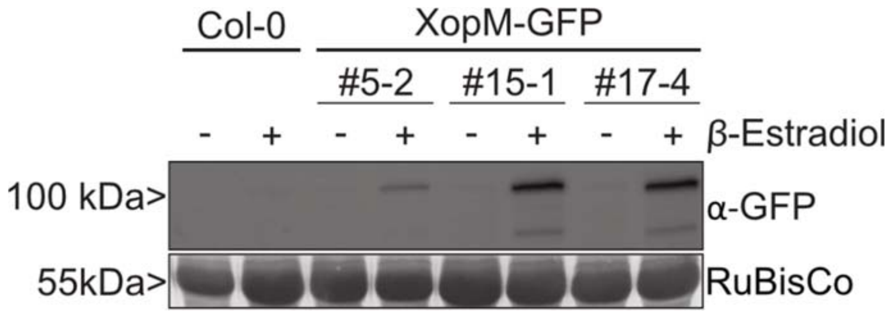


**Supplementary Figure 2: Immunoblot analysis of inducible of XopM-GFP expression in transgenic Arabidopsis lines.** Western blot analysis to confirm XopM-GFP protein expression in β-estradiol inducible transgenic *Arabidopsis* lines. Two-week-old Col-0 and XopM-GFP transgenic Arabidopsis seedlings were treated with 50 μM β-Estradiol (+) to induce XopM-GFP production or 0.1% Ethanol (-) as a control. Samples were harvested one day after induction. Western blot with anti-GFP antibody confirms the expression of XopM-GFP in the β-Estradiol induced samples. RuBisCo staining with AmidoBlack acts as a loading control.

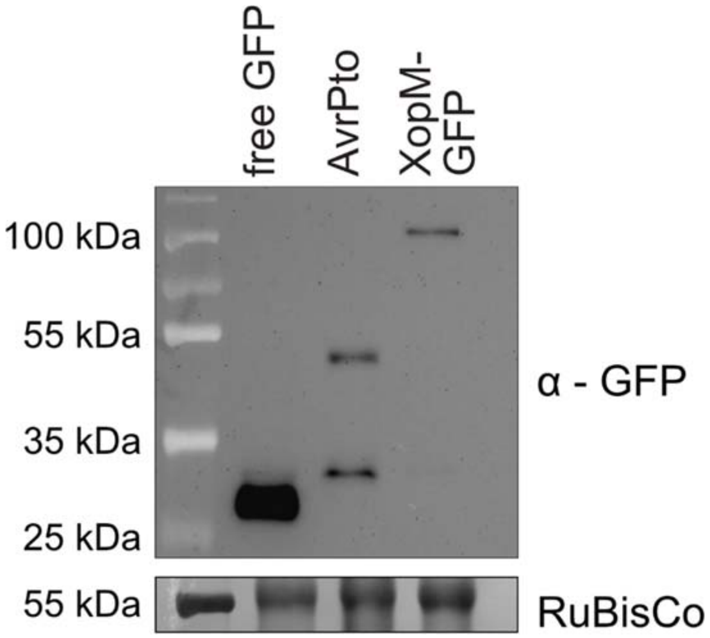


**Supplementary Figure 3. Verification of effector protein expression for ROS and POX assays.** Total protein extracts from *Agrobacterium*-infiltrated *N. benthamiana* leaves were prepared 48 hpi and protein expression was immunodetected using an anti-GFP antibody. Amido black staining of RubisCo served as loading control. Leaf discs from the biological replicates used for ROS and POX assays were pooled for the western blot analysis shown here.

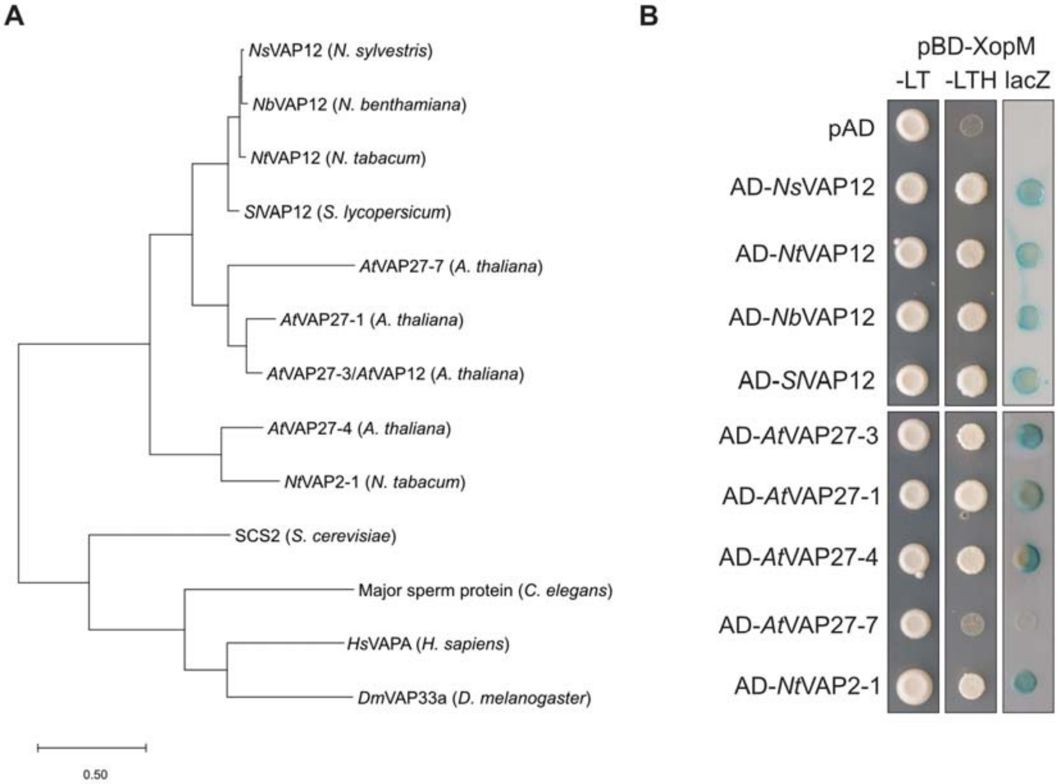


**Supplementary Figure 4: XopM interacts with several VAP isoforms.** (A) Phylogenetic tree of VAPs of plants and other model species. The tree was generated using the maximum likelihood method. Branch length give the number of substitutes per site. (B) XopM fused to the GAL4 DNA binding domain (BD) was co-expressed with VAP proteins from different plant species fused to the GAL4 activation domain (AD). The empty vector containing the AD (pAD) was used as a negative control. –LT yeast grown on selective medium lacking Leu and Trp. –LTH, yeast grown on selective medium lacking Leu, Trp and His. LacZ, activity of the lacZ reporter gene. *Ns*, *N. sylvestris*; *Nt*, *N. tabacum*; *Nb, N. benthamiana; At, A. thaliana; Sl, S. lycopersicum*. Evolutionary analysis for (A) was conducted with MEGA X (www.megasoftware.net).

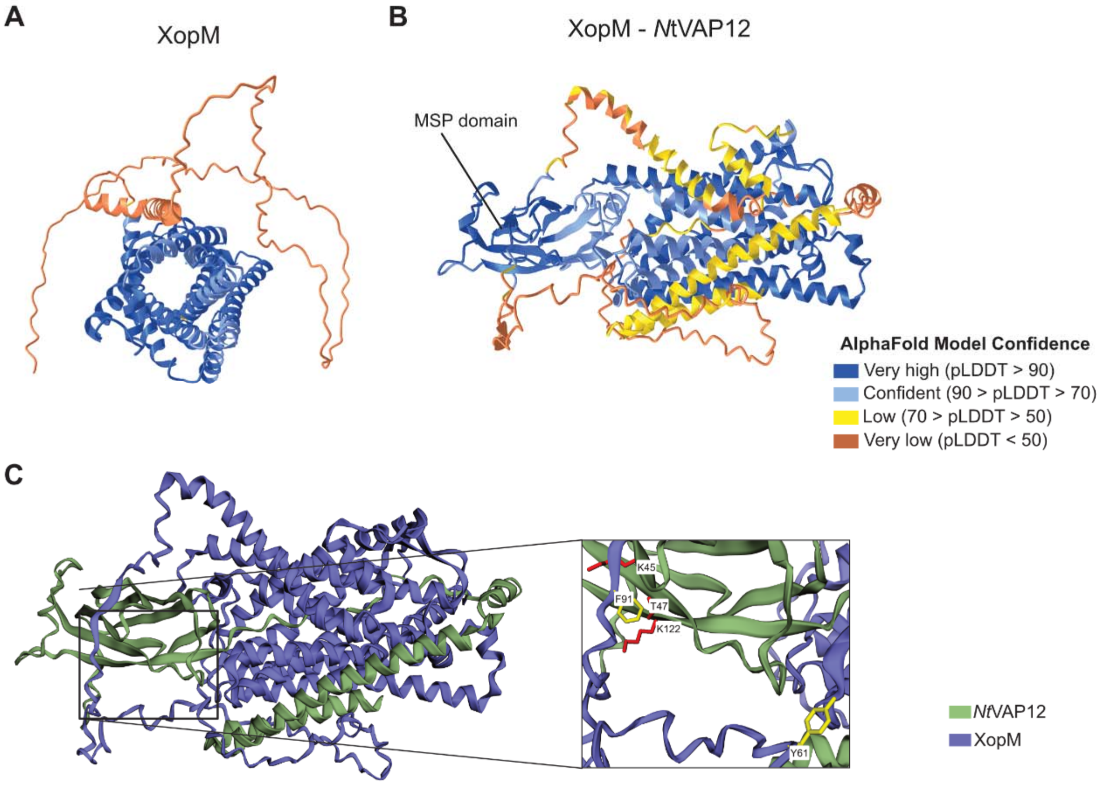


**Supplementary Figure 5. Structural modelling of XopM and XopM with *Nt*VAP12.** Cartoon representation of the predicted structures of XopM by itself and XopM with *Nt*VAP12. Structural modelling was done using AlphaFold. (A) XopM structure with high confidence levels in the C-terminal region, that has multiple alpha-helices. (B) NtVAP12's major sperm domain (MSP) made up of beta sheets is highlighted. Model confidence scores represented by colour. pLDDT, predicted local-distance difference test. Images were prepared using iCn3D. (C) Model of the XopM-*Nt*VAP12 interaction, highlighting the site of interaction. The two XopM residues (Y61/F91) essential for the interaction are highlighted in yellow. Images were prepared using EzMol.

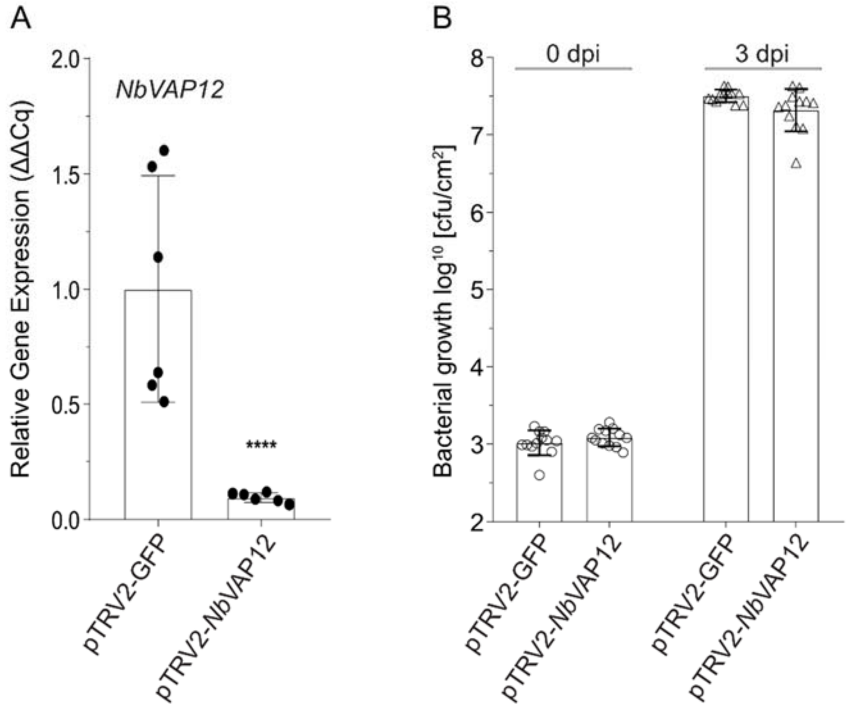


**Supplementary Figure 6: Bacterial growth of *Xcv* is not affected by reduced expression of *NbVAP12*.**

(A) Downregulation of *NbVAP12* gene expression is verified in pTRV2-*NbVAP12 roq1 N. benthamiana* plants. RNA was extracted from about four-week-old plants that were silenced two weeks prior. Quantities of relative gene expression of *NbVAP12* was measured via qRT-PCR, with *NbActin* being used as the control gene. The individual values and average of n=6 samples are plotted with the error bars displaying the standard deviation (SD). Statistically significant difference was calculated using an unpaired t-test with $p<0.0001$ indicated as ****. (B) Bacterial growth of *NbVAP12* silenced plants show no difference to non-silenced plants. Approximately five-week-old *roq1 N. benthamiana* leaves were infected with *Xcv* by pressure infiltration with a bacterial density of $1\times10^5$ CFU $mL^{-1}$. Colony forming units of the infected leaf tissue was determined on the day of infection (0 dpi) and five days post infection (5 dpi). Individual values with the bars giving the mean of six biological replicates (n=6) and two technical replicates per biological replicate. Statistical difference between the treatments on each day was calculated using a t-test with no statistical difference between treatments ($p>0.05$).

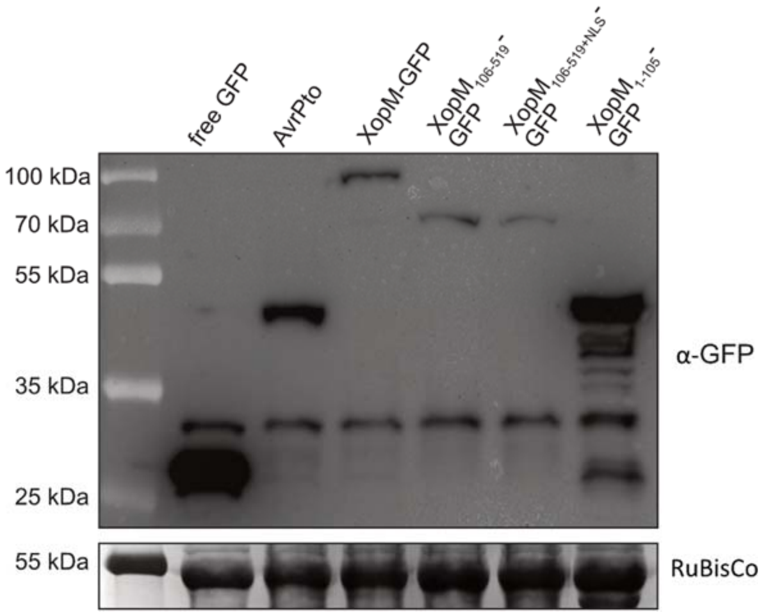


**Supplementary Figure S7. Verification of protein expression of XopM protein variants for ROS assays.** Total protein extracts from *Agrobacterium*-infiltrated *N. benthamiana* leaves were prepared 48 hpi and protein expression was immunodetected using an anti-GFP antibody. Amido black staining of RubisCo served as loading control. Leaf discs from the biological replicates used for ROS assays were pooled for the western blot analysis shown here.

Supplementary Table S1. Oligonucleotides used in the study

| | | | |
|---|---|---|---|
| CB35 | Fwd | CTTTTAAGGTCAAAGCTACCAACCCG | Mutagenesis *NtVAP* T46A |
| CB36 | Rev | CGGGTTGGTAGCTTTGACCTTAAAAG | |
| CB37 | Fwd | CTTTTAAGGTCAAAACTGCCAACCCG | Mutagenesis *NtVAP* T47A |
| CB38 | Rev | CGGGTTGGCAGTTTTGACCTTAAAAG | |
| CB39 | Fwd | GCTTTTAAGGTCAAAGCTGCCAACCCGAAAAAAATATTG | Mutagenesis *NtVAP* T46A/T47A |
| CB40 | Rev | CAATATTTTTTCGGGTTGGCAGCTTTGACCTTAAAAGC | |
| CB41 | Fwd | AGCGAGCAGGCCTACGACGCTGAAGACTC | Mutagenesis *XopM* F91A |
| CB42 | Rev | GAGTCTTCAGCGTCGTAGGCCTGCTCGCT | |
| CB43 | Fwd | CAGTTCTACGACGATGAAGACTCCTTG | Mutagenesis *XopM* A94D |
| CB44 | Rev | CAAGGAGTCTTCATCGTCGTAGAACTG | |
| CB45 | Fwd | GCAGCGAGCAGGCCTACGACGATGAAGACTCCTTGG | Mutagenesis *XopM* F91A/A94D |
| CB46 | Rev | CCAAGGAGTCTTCATCGTCGTAGGCCTGCTCGCTGC | |
| CB47 | Fwd | AAACCCAAAAAAAGAGATCGCAATGAGTAACGAGCTTCTC | *AtVAP27-3* in pGAD424 |
| CB48 | Rev | CAGGTCGACGGATCCCCGGGTCATGTCCTCTTCATAATG | |
| CB49 | Fwd | AAACCCAAAAAAAGAGATCGCAATGAGTAACGGAGGAGAG | *SlVAP12* in pGAD424 |
| CB50 | Rev | CAGGTCGACGGATCCCCGGGTTATGTCCTTTTGAGAAGATAG | |
| CB54 | Fwd | CACCAACAAAATGAGCAGTACAGAACAAG | *NtVAP12* in pENTR™/D-TOPO® |
| CB24 | Rev | GGATCAACTTTTGAGGATGTAG | |
| CB57 | Fwd | CACCAACAAAATGACGAAGATCTC | *XopM* in pENTR™/D-TOPO® |
| CB15 | Rev | GGATCCCTCAGTTCGCGTGCGAGCGCCCTCGG | |
| CB91 | Fwd | AAAGAGATCGAATTCCCGGCCATGAGTAACATCGATCTGATTG | *AtVAP27-1* in pGAD424 |
| CB92 | Rev | TAGATCTCTGCAGGTCGACGTTATGTCCTCTTCATAATGTATC | |
| CB93 | Fwd | AAAGAGATCGAATTCCCGGCCATGAGTGACGAGCTTCTC | *AtVAP27-7* in pGAD424 |
| CB94 | Rev | TAGATCTCTGCAGGTCGACGCTAAGTAGGAGATAACCAAAC | |
| CB97 | Fwd | AAAGAGATCGAATTCCCGGCCATGACTGGTACTACCAAC | *NtVAP2-1* in pGAD424 |
| CB98 | Rev | TAGATCTCTGCAGGTCGACGCTATTCTGTAGAAGGTGAAG | |
| CB109 | Fwd | GCGACAGCGGAGCTGAAACCGCCAAC | Mutagenesis *XopM* Y61A |
| CB110 | Rev | GTTGGCGGTTTCAGCTCCGCTGTCGC | |
| CB118 | Rev | GGATCCTGATGGCAAAGCCGACG | $\textit{XopM}_{1-105}$ in pENTR™/D-TOPO® |
| CB119 | Fwd | CACCAACAAAATGCTTCCCCAGGAACAC | $\textit{XopM}_{106-519}$ in pENTR™/D-TOPO® |

| | | | |
|---|---|---|---|
| CB138 | Fwd | CGACGACAAGACCCTGGAACTTCTGAGGCG | *NbVAP12* in pTRV2 |
| CB139 | Rev | GAGGAGAAGAGCCCTAGTTGCAACTTCTTTTG | |
| CB182 | Fwd | AGATCGAATTCCCGGGAATGACCGGCGTTGGCG | *AtVAP27-4* in pGAD424 |
| CB183 | Rev | CTCTGCAGGTCGACGTTATGTGGGAGAAGCTAAGGTAAGTTTT | |
| CB184 | Fwd | AAACCCAAAAAAAGAGATCGCAATGAGCAGTGCAGAACAAGAACTTCTCAACATCGATCC | *NbVAP12* in pGAD424 |
| CB185 | Rev | CGACGGATCCCCGGGTCAATTTTTGAGGATGTAGCCCAAAGCTACACC | |
| CB188 | Fwd | aaacccaaaaaaagagatcgaaCTTCTCGACATCAATCCTC | $MSP_{NtVAP12}$ in pGAD |
| CB189 | Rev | cgacggatccccgggGGGAAGATAAATCACTCTC | |
| CB193 | Fwd | CACCATGccaaaaaaaaaaagaaaagttCTTCCCCAGGAACACG | NLS-XopM inpENTR™/D-TOPO® |
| FB918 | Fwd | GCGGATCCATGACGAAGATCTCTTCAGCCAG | *Xcv ΔXopM* N-terminal fragment in pOK |
| FB919 | Rev | GGCGACCTTGAGCTCACTGGAGGGTGGTGACTCCATG | |
| FB920 | Fwd | CATGGAGTCACCACCCTCCAGTGAGCTCAAGGTCGCC | *Xcv ΔXopM* C-terminal fragment in pOK |
| FB921 | Rev | GTCGACTCACTCAGTTCGCGTGCGAGCGC | |
| *Nb*VAP12 | Fwd | ACATCACAGCAACAGACTTCGT | qPCR |
| | Rev | AACATGAAGGAAGCACTACCA | |
| *Nb*Actin | Fwd | GCCAACAGAGAGAAGATGACCCAGA | qPCR |
| | Rev | ACACCATCACCAGAGTCCAACACAAT | |
